# Lipid levels correlate with neuronal and dopaminergic markers during the differentiation of SH-SY5Y cells

**DOI:** 10.1101/2023.09.01.555964

**Authors:** Frederik Ravnkilde Marlet, Sonia Sanz Muñoz, Nefeli Sotiraki, Jannik Nicklas Eliasen, Jakob Paul Woessmann, Jan Weicher, Jesper Elmsted Dreier, Erwin Schoof, Kristi A Kohlmeier, Kenji Maeda, Céline Galvagnion

**Affiliations:** Department of Drug Design and Pharmacology, Faculty of Health and Medical Sciences, University of Copenhagen, 2100, Copenhagen, Denmark; Cell Death and Metabolism Unit, Center for Autophagy, Recycling and Disease, Danish Cancer Society Research Center, Copenhagen, Denmark; Department of Biotechnology and Biomedicine, Technical University of Denmark, Lyngby 2800, Denmark

## Abstract

Parkinson’s Disease (PD) is characterized by the loss of dopaminergic neurons and the deposition in the remaining cells of protein inclusions called Lewy Bodies (LBs). LBs are heterogeneous structures composed of protein and lipid molecules and their main constituent is the presynaptic protein α-synuclein. SH-SY5Y cells are neuroblastoma cells commonly used to model PD because they express dopaminergic markers and α-synuclein and they can be differentiated into neuronal cells using established protocols. Despite increasing evidence pointing towards a role of lipids in the initiation of PD, limited knowledge is available on the lipidome of undifferentiated and differentiated SH-SY5Y cells. In this study, we show that the levels and chemical properties of negatively charged phospholipids, diacyl glycerol and sphingolipids are specifically altered along the differentiation process of SH-SY5Y cells and that the levels of these lipids’ species correlate with those of dopaminergic and neuronal markers. These results are supported by proteomic data showing that the main biological processes affected by the differentiation of SH-SY5Y cells are lipid metabolism and processes associated with neuron maturation. Finally, our results show that electrophysiological activity can be detected in differentiated SH-SY5Y cells at a stage where most of the lipid changes have reached their maximal value. These results provide the first complete and quantitative characterisation of the changes in lipidome associated with the differentiation of SH-SY5Y cells into more neuronal and dopaminergic-like phenotype and serve as a basis for further characterisation of lipid disruptions in association with PD and its risk factors in this dopaminergic-like neuronal cell model.

## Introduction

Parkinson’s Disease (PD) is a neurological disorder characterised by two pathological hallmarks, i.e. the loss of dopaminergic neurons in the substantia nigra pars compacta and the deposition of protein inclusions called Lewy Bodies (LBs) in the remaining neurons^1^. LBs are made of proteins, lipids, and organelles and their main constituent is the small pre-synaptic protein, alpha-synuclein (αS)^2,3^. αS is an intrinsically disordered protein that can adopt an α-helical structure upon binding to lipid vesicles^4–6^. This protein-membrane interaction is proposed to play a role in its biological function, i.e. the fusion of synaptic vesicles^7^, but also to be responsible for the early events leading to the formation of amyloid fibrils^8–12^. In fact, the lipid composition of membranes was found to modulate both the thermodynamics of αS membrane binding and the kinetics of αS amyloid fibrils formation^13^. Moreover, lipid molecules do not only act as a trigger of αS aggregation by providing a surface in the form of a membrane on which the protein can start aggregating but they also act as reagents by co-assembling with αS into lipid-protein amyloid fibrils^14,15^. Indeed, we found that αS can bind to vesicles made with negatively charged phospholipids but can only co-assemble with those characterised by short chain fatty acids into amyloid fibrils^13,15^.

Increasing number of studies point towards a link between changes in lipid levels and/or properties and PD and αS pathology^16,17^. Indeed, analyses of post-mortem brain samples, patients’ fibroblasts and body fluids have shown that the levels of specific sphingolipids were disrupted in association with both the sporadic and familial form of the disease^17^. In particular, the levels of short chain sphingolipids were found to be increased in the brain of patients with PD and in the fibroblasts of PD patients with *GBA* mutations^18,19^. *GBA* encodes Glucocerebrosidase (GCase), a lysosomal glycoprotein involved in the hydrolysis of glucosylceramide into ceramide and glucose^20^. Mutations in the *GBA* gene is the largest genetic risk factor of the disease and have been associated with an earlier onset of the disease^21–23^.

SH-SY5Y neuroblastoma cell line is commonly used to model PD and αS pathology and the disease risk factors via e.g. genetic modification (overexpression of αS WT and disease variants, knock-down of gene associated with PD), exposure to neurotoxic agents (1-methyl-4-phenylpyridinium, 6-hydroxydopamine), enzyme inhibitors or pre-formed fibrils and/or *ex-vivo* oligomers^24–38^. SH-SY5Y cells are derived from the parental line SK-N-SH cells^39^, which is a mixture of 3 different cell types: the neuroblastic type (N-type), which contains small cell bodies with neurite-like processes, the substrate-adherent type (S-type), which has flat and larger cell bodies, and both of them can undergo transdifferentiation through an intermediate type (I-type)^40^. This bidirectional interconversion affects not only the cell morphology, but also changes the biochemical properties of the cells^41^. Indeed, Ferlemann et al. profiled the different subtypes of SH-SY5Y cells using surface markers and showed that the treatment of the cells with the small molecule bone morphogenetic protein 4 (BMP4) led to the reduction of the population of S-type and the enhancement of the expression of the N-type cluster^42^. In order to obtain a more homogeneous population of cells with a more neuronal and dopaminergic-like phenotype, SH-SY5Y cells are commonly differentiated using retinoic acid (RA) alone, or in combination with other agents such as 12-O-Tetradecanoylphorbol-13-acetate (TPA), brain-derived neurotrophic factor (BDNF) or dibutyryl cyclic AMP^36,43,44^. The effects of these treatments on the phenotype of SH-SY5Y cells have been investigated only in a limited number of publications by assessing the levels of dopaminergic and mature neuronal markers using qPCR, Western Blot (WB), immunocytochemistry (ICC) and transcriptional profiling ^28,32,45–51^. Undifferentiated cells express dopaminergic neuronal markers including tyrosine hydroxylase (TH), dopamine transporter (DAT) and dopamine receptor 2 and 3 (D2R and D3R), as well as immature neuronal markers such as nestin ^47,52–54^. RA-induced differentiation was found to increase neurite length^49,51,55,56^ and to either have no effect or to increase the levels of dopaminergic markers in SH-SY5Y cells^24,28,46–51,57,58^. The combined RA/TPA-induced differentiation of SH-SY5Y cells was, however, found to systematically lead to a significant increase in the levels of TH, DAT, D2R and D3R as assessed by ICC and WB^28,32,48,59^. Finally, the differentiation of SH-SY5Y cells was found to inconsistently increase the levels of endogenous αS^31,32,34,60^.

Although SH-SY5Y cells have been used to model PD for the past decades, no information on the lipidome of the undifferentiated and differentiated cells is yet available. Our study aims at filling this knowledge gap and reports the change in the lipidome of SH-SY5Y cells associated with their differentiation into more neuronal and dopaminergic-like cells with a higher electrophysiological activity. Our shotgun lipidomic analyses showed that the differentiation of SH-SY5Y cells leads to an increase in the levels of Sphingolipids (SL) (Hexosyl Ceramide (HexCer), Sphingomyelin (SM), Ceramide (Cer)) and Diacylglycerides (DAG) and a decrease in those of negatively charged phospholipids Phosphatidylinositol (PI), Phosphatidylserine (PS) and Phosphatidic Acid (PA). The differentiation of the SH-SY5Y cells was found to also change the chain distribution of the phospholipids, ether phospholipids, DAG and SL. Among the species affected by the differentiation process, we could identify changes in the levels of specific species common for all sphingolipids, i.e. the increase in the levels of the 36:1 species and the decrease in those of 40:1 species for SM, HexCer and Cer. Moreover, we observed that the differentiation of SH-SY5Y cells led to the increase in β-III tubulin positive cells, TH levels and neurite length and that the levels of 36:1 and 40:1 SL correlate positively and negatively, respectively, with those of TH and/or the neurite length. Finally, we found that SH-SY5Y cells starts showing electrophysiological activity after the RA treatment which correspond to the time where most changes in lipid levels have reached their maximum or minimum values. All together, these results provide a complete and quantitative characterisation of the changes in lipidome associated with the differentiation of SH-SY5Y cells into a more neuronal and dopaminergic-like phenotype. Our study serves as a basis for the use of this dopaminergic-like neuronal cell model for the characterisation of lipid disruptions in association with PD and its risk factors.

## Results

### The differentiation of SH-SY5Y cells leads to significant changes in the levels and chain properties of glycerophospholipids, diacylglyceride and sphingolipids

We used shotgun lipidomics to quantitively characterise the effect of the differentiation of SH-SY5Y cells on their lipidome. The cells were differentiated as four independent preparations by sequential treatments with BMP4, RA and TPA (BRT treatments) in the presence and absence of FBS. The following three conditions were considered: (i) B/R+/T+, (ii) B/R+/T− and (iii) B/R−/T−, which correspond to the successive treatments of the cells using BMP4 (5% FBS), RA (1% FBS) and, TPA (1% FBS) for B/R+/T+, or BMP4 (5% FBS), RA (1% FBS) and TPA (no FBS) for B/R+/T−, or BMP4 (5% FBS), RA (no FBS) and TPA (no FBS) for B/R−/T−. We quantified the lipidome of the cells in the undifferentiated state, i.e. after 6 days in maintenance media (“UD”), and along the differentiation process, i.e. after a 6 day treatment with BMP4 (“BMP4”), or after a subsequent 6 day treatment with RA in the presence and absence of FBS (“B/R+” and “B/R-“, respectively) or after a final 6 day treatment with TPA in the presence and absence of FBS (“B/R+/T+”, “B/R+/T−” and “B/R−/T−”, respectively). The shotgun lipidomic analyses of SH-SY5Y cells at different time points allowed us to quantify more than 430 unique lipid species belonging to 25 lipid classes. Data were converted from picomol to percentage of total identified lipids per sample for each lipid in order to allow the direct comparison of samples with different total lipid amounts. Figure 1a shows a principal-component analysis (PCA) of the shotgun lipidomic analyses of SH-SY5Y cells at different time points along the differentiation process. The PCA plot shows that the four replicates clustered together, demonstrating excellent reproducibility from one differentiation to another. We observed that the BMP4 treated cells cluster more together than the UD cells, indicating that the BMP4 treatment alone helps setting the cells to a more homogeneous starting point for differentiation. We also observed that B/R+/T− and the B/R−/T− treated cell preparations cluster more together than the B/R+ and B/R- cell preparations, suggesting that these treatments are likely to render the cell population more homogeneous.

**Figure 1:**
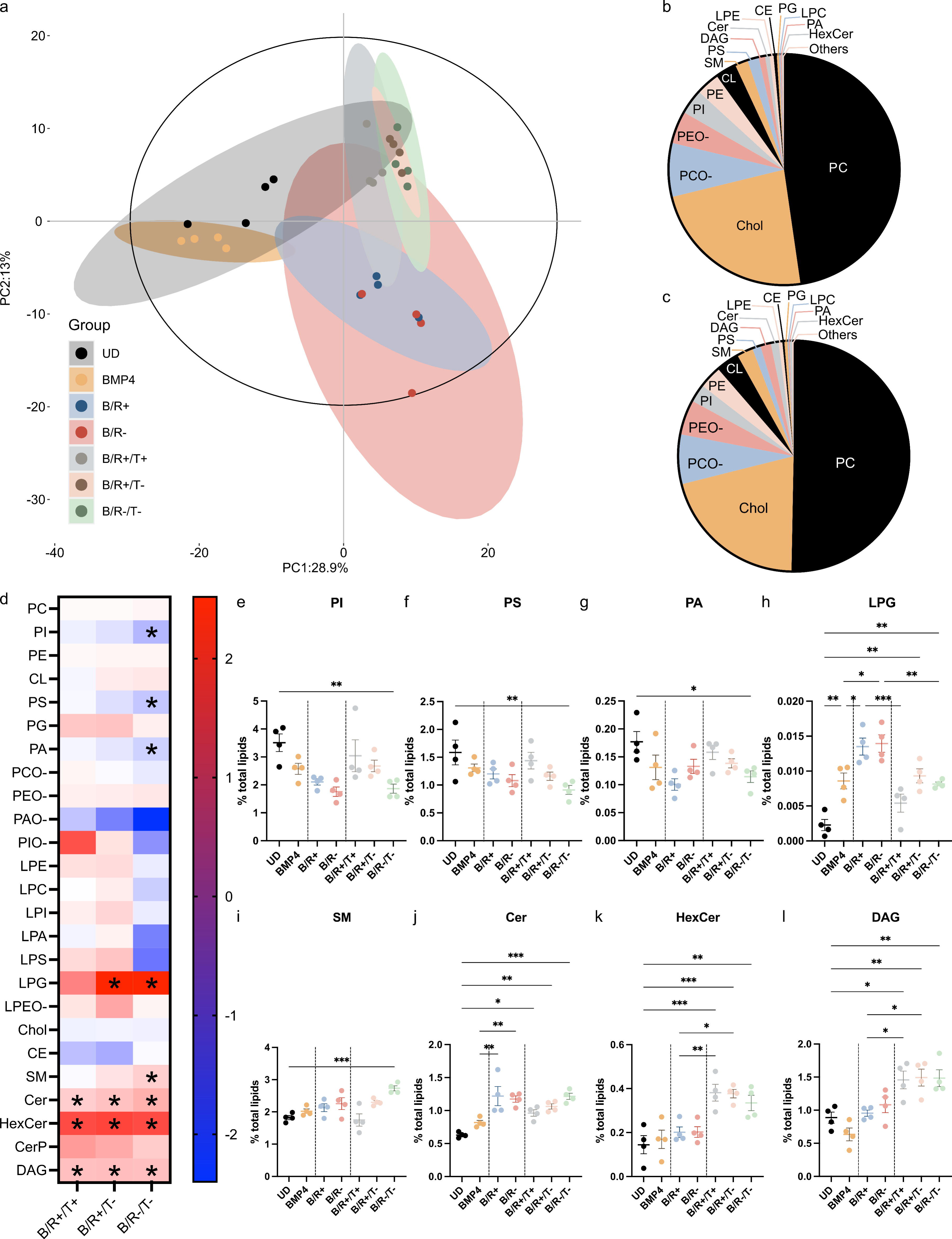
The differentiation of SH-SY5Y cells leads to significant changes in the levels of negatively charged phospholipids, sphingolipids and DAG. a. PCA of the lipidome of SH-SY5Y cells at different stages of the differentiation process, where the filled out ellipses correspond the 95% confidence intervals of each group and the black outlined ellipse is the Hotelling t-squared distribution of all groups and replicates. b, c. Average distribution of the 18 most abundant lipids in undifferentiated (b, n = 4) and differentiated (c, n= 12 (average B/R+T/+, B/R+/T−, B/R−/T−)) SH-SY5Y cells displayed as parts of the whole. d. Heat map showing the log2 of the fold-change (log2(FC)) in the levels of lipid classes upon BRT differentiation. Significant changes (p<0.033, p<0.01 and p<0.001) in log2(FC) are all indicated by one asterisk for clarity. e-l. Changes in the levels of lipid classes along the differentiation process. Lipid classes whose levels are significantly affected by at least one BRT treatment and that have been detected in all UD and BRT replicates are shown. One-way ANOVA to compare means of UD, BMP4, B/R+, B/R-, B/R+/T+, B/R+/T− and B/R−/T−, *p<0.033, **p<0.002, ***p<0.001 (see Table S1 for p-values and statistical analysis details). Comparison shown: B/R+/T+, B/R+/T− and B/R−/T− vs UD (d), all vs UD, B/R+ and B/R- vs BMP4, B/R+/T+, B/R+/T− and B/R−/T− vs their respective B/R(+/-) (e-l). Data shown as mean ± SEM and derive from 4 independent differentiations. Abbreviations of lipid classes are defined by Lipid Maps classification.

The most abundant lipids in the undifferentiated SH-SY5Y cells were Phosphatidylcholine (PC) (48%), Cholesterol (23%), ether-linked Phosphatidylcholine (PCO-) (7.6%), ether-linked Phosphatidylethanolamine (PEO-) (4.5%), PI (3.5%), Phosphatidylethanolamine (PE) (3.3%), Cardiolipin (CL) (3.0%), SM (1.8%), and PS (1.6%) (Figure 1b).

The differentiation of SH-SY5Y cells led to the significant decrease in the levels of PI, PS and PA as well as the significant increase in the levels of lyso Phosphatidylglycerol (LPG), SM, Cer, HexCer and DAG consistently for one or more BRT conditions (Figure 1d). We observed that the decrease of PS and the increase of SM and Cer were more pronounced for the conditions without serum, i.e. B/R+/T− and/or B/R−/T−, compared to B/R+/T+ (Figure 1d and S1a). The levels of these lipids varied along the differentiation process via different patterns. The changes in the levels of PI, PS, PA and SM occurred progressively until reaching significance at the end of the BRT treatment whereas those of Cer, HexCer and DAG occurred by steps and attain their maximal levels after the RA (Cer) or TPA treatment (HexCer, DAG) (Figure 1e-g, i-l). Finally, the levels of LPG first increased after BMP4 treatment and then again after RA treatment, when they reached their highest value, and finally decreased after the TPA treatment (Figure 1h). We also identified a group of glycerophospho- and lyso-glycerophospholipids whose levels were significantly altered along the differentiation process even though their levels at the end of the BRT treatment were the same as those in the undifferentiated cells. In particular, we observed that the levels of PC, PCO-, PAO- and PIO- were all significantly increased after the RA treatment and then decreased after the TPA treatment in the absence and/or presence of FBS (PC (B/R+/T+), PAO- and PIO- (B/R−/T−), PCO- (B/R+/T+, B/R+/T−, B/R−/T−) (Figure S2). We note that BMP4 pre-treatment led to a significantly more pronounced increase in SM levels upon differentiation when comparing B/R+/T− vs R+/T− SH-SY5Y cells (Figure S3).

We then investigated the effect of the differentiation of SH-SY5Y cells on the levels of the lipid species, i.e. on the chain distribution of all lipid classes. Data were converted from percentage of total identified lipids to percentage of lipid class (e.g. percentage of PC) for each lipid in order to allow the direct comparison of samples with different total lipid class (e.g. different % of PI in undifferentiated vs differentiated cells). Figure 2, Table 1 and Figure S4-13 summarise the changes in the levels of the most abundant lipid species (level higher than 5% of the total lipid class) detected in the undifferentiated and/or differentiated cells. Although the chain distribution varied from one glycerophospholipid to another and from glycerophospholipids to DAG, we could identify subspecies whose levels varied significantly in the same direction across the different glycerophospholipids and DAG. The levels of 36:2, 36:3, 38:3, 40:3 and/or 40:4 were systematically increased in the differentiated cells for at least one BRT condition for PC, PCO-, PEO-, PI, PE, PS, PG, PA and/or DAG (Table 1). On the contrary, the differentiation of SH-SY5Y cells led to a decrease in the levels of 36:1 subspecies for PC, PE and PS (Table 1). Finally, the levels of a group of species were increased for some lipids and decreased for others in BRT treated SH-SY5Y cells compared to UD cells. In particular, the levels of 32:1 were increased for PC, PCO- but decreased for PA and DAG, those of 34:1 were decreased for PC, PE, PG and DAG and increased for PCO-, PS and those of 34:2 were increased for PC, PCO-, PEO-, PG and PA when comparing BRT vs UD SH-SY5Y cells (Table 1). We found that the significant increase in the levels of these lipids upon differentiation were more pronounced in the conditions where no serum was used during the RA and/or TPA treatment, i.e. when comparing B/R+/T− and B/R−/T− treated cells to B/R+/T+ (Table 1 and Figure S1). We noted that BMP4 pre-treatment led to significantly more pronounced increase in the levels of the 36:2 species of PC, PE and PS, PI 38:3 and PCO- 34:2 and decrease in the levels of PI 38:4 upon differentiation of SH-SY5Y cells when comparing B/R+/T− vs R+/T− cells (Figure S3). The observed changes in species levels upon BRT differentiation occurred either progressively along the differentiation process until they reached significance at the end of the BRT treatment or by step after BMP4, RA and/or TPA treatments (Figure S4-13), suggesting that the combined BRT treatment was required for all glycerophospholipids and DAG species to reach their maximal/minimal values. The chain of the sphingolipids (SL), i.e. Cer, SM and HexCer, were also significantly altered by the differentiation but in a more systematic way (Figure 2 and S13). We observed that the levels of 36:1 were increased for all SL in BRT-induced differentiated cells compared to undifferentiated cells and that such an increase made the 36:1 SL species changed from one of the minor species in undifferentiated cells to one of the most abundant species after BRT-induced differentiation. On the contrary, the levels of 40:1 (SM, Cer, HexCer) or 42:2 (SM) were decreased in the differentiated SH- SY5Y cells compared to the undifferentiated cells (Figure 2b-h).

**Figure 2:**
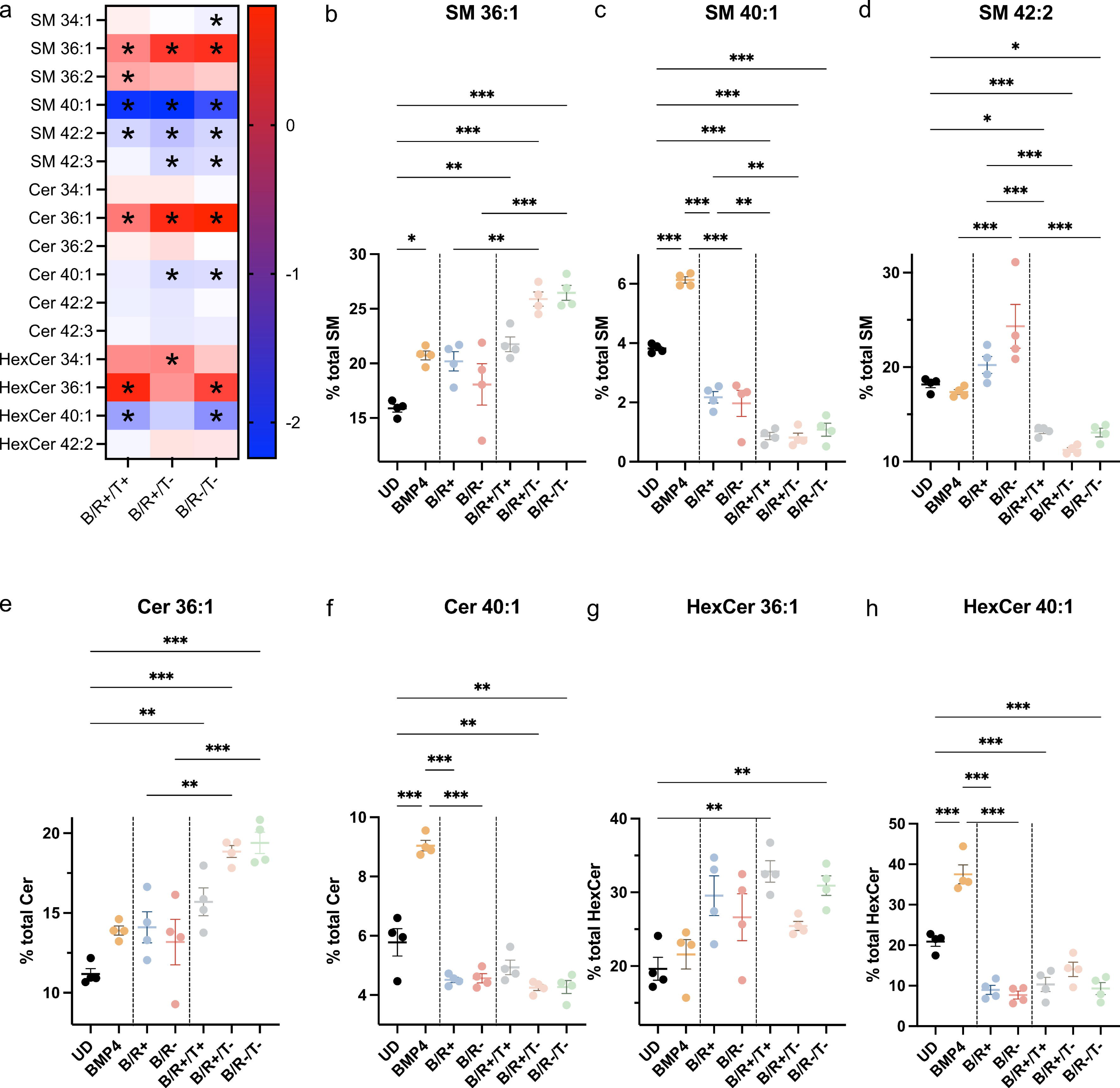
The differentiation of SH-SY5Y cells leads to significant changes in the chain distribution of sphingolipids. a. Heat map showing the log2(FC) in the levels of the most abundant species of sphingolipids upon BRT differentiation. Significant changes (p<0.033, p<0.01 and p<0.001) in log2(FC) are all indicated by one asterisk for clarity. b-h. Change in the percentage of the most abundant sphingolipids (in mole percentage of the class) whose levels are significantly increased or decreased by the BRT differentiation along the differentiation process. One-way ANOVA to compare means of UD, BMP4, B/R+, B/R-, B/R+/T+, B/R+/T− and B/R−/T− (a-h), *p<0.033, **p<0.002, ***p<0.001 (see Table S1 for p-values and statistical analysis details). Comparison shown: B/R+/T+, B/R+/T− and B/R−/T− vs UD (a), all vs UD, B/R+ and B/R- vs BMP4, B/R+/T+, B/R+/T− and B/R−/T− vs their respective B/R(+/-) (b-h). Data shown as mean ± SEM and derive from 4 independent differentiations.

**Table 1:**
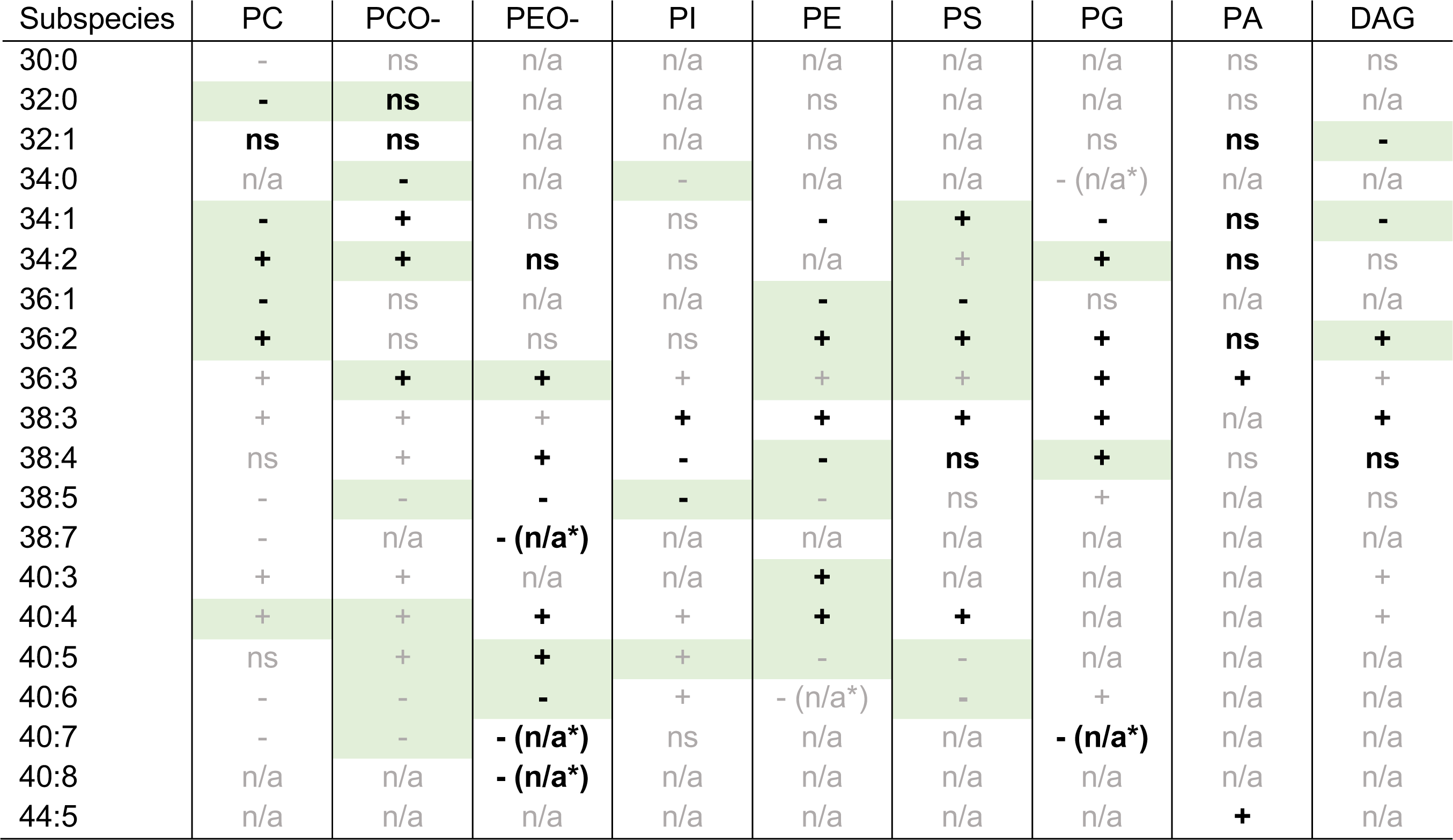
Significant changes in the levels of DAG, phospholipid and ether phospholipid subspecies associated with the BRT differentiation of SH-SY5Y cells. One-way ANOVA to compare the means of UD, B/R+, B/R-, B/R+/T+, B/R+T/-, B/R−/T− see Figure S4-12 and Table S1. ns, not significant, +/-: significant increased/decreased in BRT compared to UD SH-SY5Y cells; n/a: not detectable in replicates from UD and BRT, - (n/a*): main species detected in UD but not in at least one replicate of the BRT SH-SY5Y cells. For each class, the changes in the levels of the main subspecies are shown in black and bold. The subspecies whose levels are significantly higher/lower in B/R+/T− and/or B/R−/T− compared to B/R+/T+ are highlighted in green (One-way ANOVA to compare the means of the three BRT conditions, see Figure S1a).

| Subspecies | PC | PCO- | PEO- | PI | PE | PS | PG | PA | DAG |
| --- | --- | --- | --- | --- | --- | --- | --- | --- | --- |
| 30:0 | - | ns | n/a | n/a | n/a | n/a | n/a | ns | ns |
| 32:0 | - | **ns** | n/a | n/a | ns | n/a | n/a | ns | n/a |
| 32:1 | **ns** | **ns** | n/a | n/a | ns | n/a | ns | **ns** | - |
| 34:0 | n/a | - | n/a | - | n/a | n/a | - (n/a*) | n/a | n/a |
| 34:1 | - | + | ns | ns | - | + | - | **ns** | - |
| 34:2 | + | + | **ns** | ns | n/a | + | + | **ns** | ns |
| 36:1 | - | ns | n/a | n/a | - | - | ns | n/a | n/a |
| 36:2 | + | ns | ns | ns | + | + | + | **ns** | + |
| 36:3 | + | + | + | + | + | + | + | + | + |
| 38:3 | + | + | + | + | + | + | + | n/a | + |
| 38:4 | ns | + | + | - | - | **ns** | + | ns | **ns** |
| 38:5 | - | - | - | - | - | ns | + | n/a | ns |
| 38:7 | - | n/a | - (n/a*) | n/a | n/a | n/a | n/a | n/a | n/a |
| 40:3 | + | + | n/a | n/a | + | n/a | n/a | n/a | + |
| 40:4 | + | + | + | + | + | + | n/a | n/a | + |
| 40:5 | ns | + | + | + | - | - | n/a | n/a | n/a |
| 40:6 | - | - | - | + | - (n/a*) | - | + | n/a | n/a |
| 40:7 | - | - | - (n/a*) | ns | n/a | n/a | - (n/a*) | n/a | n/a |
| 40:8 | n/a | n/a | - (n/a*) | n/a | n/a | n/a | n/a | n/a | n/a |
| 44:5 | n/a | n/a | n/a | n/a | n/a | n/a | n/a | + | n/a |

The changes in the level of these SL species were found to occur progressively during the differentiation process until they reached their maximal levels (HexCer 36:1, Cer 36:1) or in steps: after BMP4 for Cer 40:1, after BMP4 and then after TPA for SM 36:1, after RA and then after TPA for SM 40:1, after RA for HexCer 40:1 and after TPA for SM 36:2 and 42:2 and HexCer 34:1 (Figure 2b-h and S13). These results reinforce our conclusion that the combined BRT treatment is required for the lipid levels to reach their maximal/minimal value in the differentiated SH-SY5Y cells. The observed increases in the levels of 36:1 species upon the differentiation of SH-SY5Y cells were specific to SL as the levels of 36:1 species for PL were found to either not be affected (PG) or be decreased (PE, PS, PC) when comparing BRT to UD cells. The comparison of our MS2/MS1 lipidomic analyses suggests that most of the 36:1 and 40:1 subspecies correspond to 18:1/18:0 and 18:1/22:0, respectively.

We note that BMP4 pre-treatment led to significantly more pronounced decrease in the levels of SM 34:1 upon differentiation of SH-SY5Y cells when comparing B/R+/T− vs R+/T− cells (Figure S3). Moreover, the increase in the levels of the 36:1 subspecies for SM and Cer was found to be significantly more pronounced for the conditions without serum, i.e. B/R+/T− and/or B/R−/T− compared to B/R+/T+ (Figure 1d and S1a).

Altogether, our shotgun lipidomic analyses of undifferentiated and BRT-induced differentiated SH- SY5Y cells show that both the levels and the chain distribution of DAG, PI, PS, PA, Cer, SM and HexCer are changed along the differentiation process. We observed that pre-treatment of the cells with BMP4 and the removal of serum during RA and/or TPA treatment led to more pronounced changes in the levels of the subspecies affected by the differentiation process. Although the levels of the most abundant DAG, PI, PS and PA subspecies were affected in different ways (increased or decreased or non-significant changes), we observed a systematic increase in the levels of 36:1 species and decrease in those of 40:1 species across all SL. We then investigated whether these lipid changes were correlated with neuronal and dopaminergic markers.

### The levels of 36:1 and 40:1 SL subspecies correlate with those of TH and neurite length

We characterised the extent by which SH-SY5Y cells became more dopaminergic and more neuronal along the process of differentiation using WB analyses for αS, TH and D3R, Image J (NeuronJ) neurite tracing and immune staining for β-III tubulin and Ki67. Our WB analyses show that the levels of TH and D3R, but not those of αS, were significantly increased in the BRT treated SH- SY5Y cells compared to the undifferentiated cells (Figure 3a-c). We observed that the BMP4 treatment alone did not affect the levels of TH, D3R and αS but a further RA treatment led to the significant increase in the level of both αS and TH when comparing BR-treated cells to BMP4 treated cells (Figure 3a-c). A subsequent treatment with TPA led to no further increase in the levels of TH but to a decrease of the levels of αS to a value similar to that in undifferentiated cells (Figure 3a-c). The neurite length analysis shows that undifferentiated and BMP4 treated SH-SY5Y cells have neuronal extension of ca. 25 µm (20-28 µm) (Figure 3d, DIV - 6 to 0, black and orange data points). After BMP4 treatment, incubation of the cells with RA then TPA led to a progressive significant increase in neurite length from ca. 25 µm at day 0 to 45 µm after 6 days in RA (Figure 3d, DIV 0 to 6, blue (B/R+) and dark pink (B/R-) data points) and to 70-130 µm after 6 additional days in TPA (Figure 3d, DIV 6 to 12, grey (B/R+/T+), light pink (B/R+/T−) and green (B/R−/T−) data points). This observed increase in neurite length upon differentiation was found to be much more pronounced for differentiated SH-SY5Y cells pre-treated with BMP4 as the neurite length of these cells were on average 1.3 times longer than those of un-pre-treated cells (Figure S3). Finally, our results show that the concentration of FBS affects the division rate and the neurite length of the differentiated SH- SY5Y cells (Figure 3d). Indeed, we found that SH-SY5Y cells divide two times slower when differentiated in the absence of serum (Figure S1b) and that B/R−/T− differentiated SH-SY5Y cells form neuronal extensions 1.4 times longer than those of B/R+/T− differentiated SH-SY5Y cells and 2.3 times longer than those of B/R+/T+ differentiated cells (Figure 3d). We however note that the cells became more fragile and would adhere less when the serum was removed from the culture media starting at the RA treatment, i.e. for the B/R- and B/R−/T− conditions.

**Figure 3:**
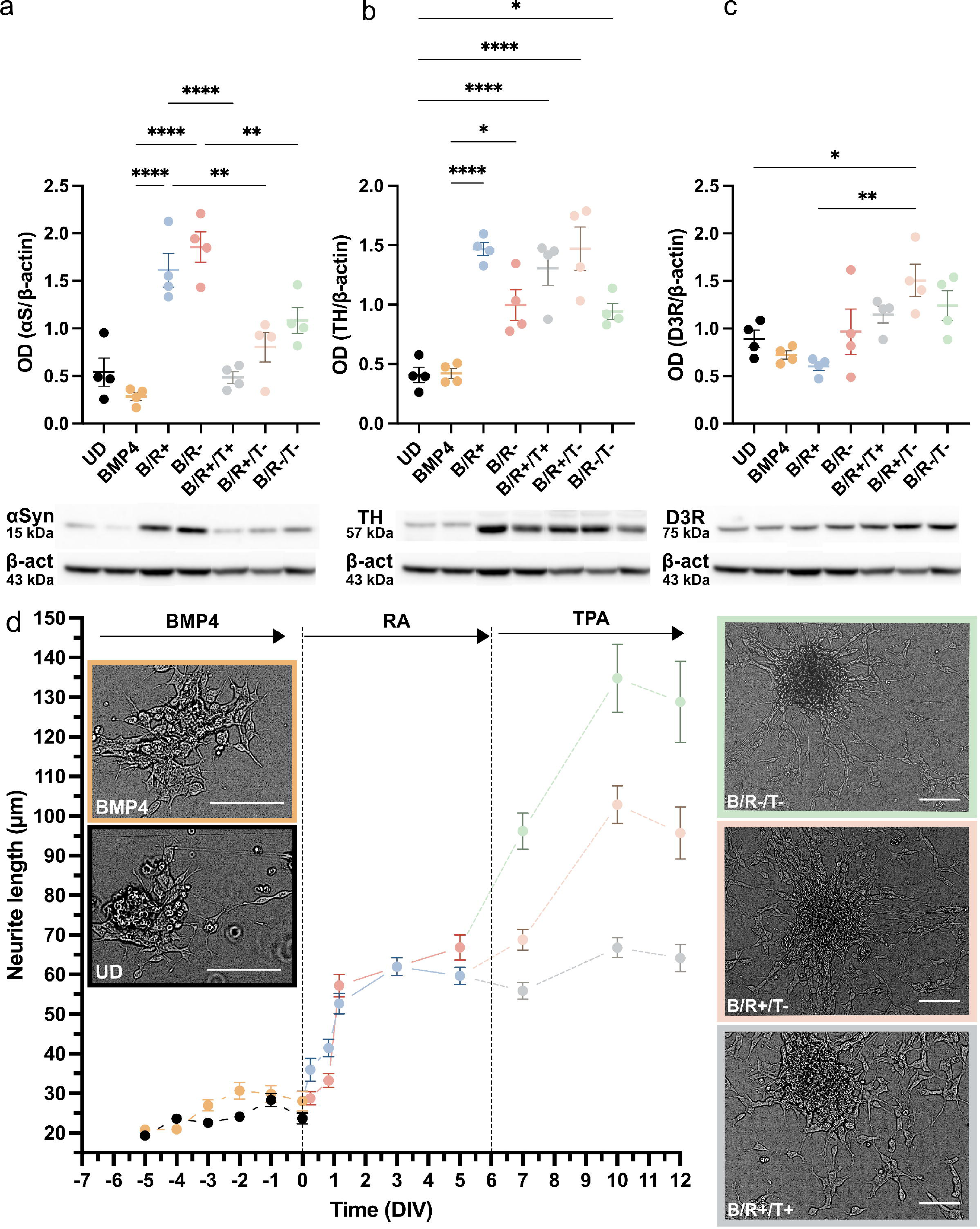
The BRT differentiation of SH-SY5Y cells leads to a significant increase in neurite length and TH levels. a-c. Semi quantitative analysis of western blot band intensity of (a) αS, (b) TH and (c) D3R normalized to β-actin. One-way ANOVA to compare means of UD, BMP4, B/R+, B/R-, B/R+/T+, B/R+/T− and B/R−/T− (a-h), *p<0.033, **p<0.002, ***p<0.001 (see Table S1 for p-values and statistical analysis details). Comparison shown: all vs UD, B/R+ and B/R- vs BMP4, B/R+/T+, B/R+/T− and B/R−/T− vs their respective B/R(+/-). Data shown as mean ± SEM and derive from 4 independent differentiations. (d) Neurite length quantification and representative bright field images at different time points of the differentiation. Scale bar = 100 µm. Data is shown as mean ± SEM derived from 100-125 individual neurite length measurements per image for a total of two images per condition per timepoint.

In order to further characterise the neuronal phenotype of SH-SY5Y cells along the BRT-induced differentiation, we stained the cells for β-III tubulin, a neuronal marker, and Ki67, a marker of cell proliferation^61^. We observed that ca. 90% and ca. 60% of the undifferentiated SH-SY5Y cells express β-III tubulin and Ki67, respectively (Figure 4). BMP4 treatment was found not to affect the number of β-III tubulin positive cells but to lead to a significant decrease in that of Ki67 positive cells (Figure 4). A further treatment with RA also did not affect the number of β-III tubulin positive cells but led to a further significant decrease in that of Ki67 positive cells. A subsequent treatment with TPA led to a significant increase in the levels of β-III tubulin positive cells compared to UD and BMP4 treated cells and an increase in the number of Ki67 positive cells to a value similar to that in BMP4 treated cells.

**Figure 4:**
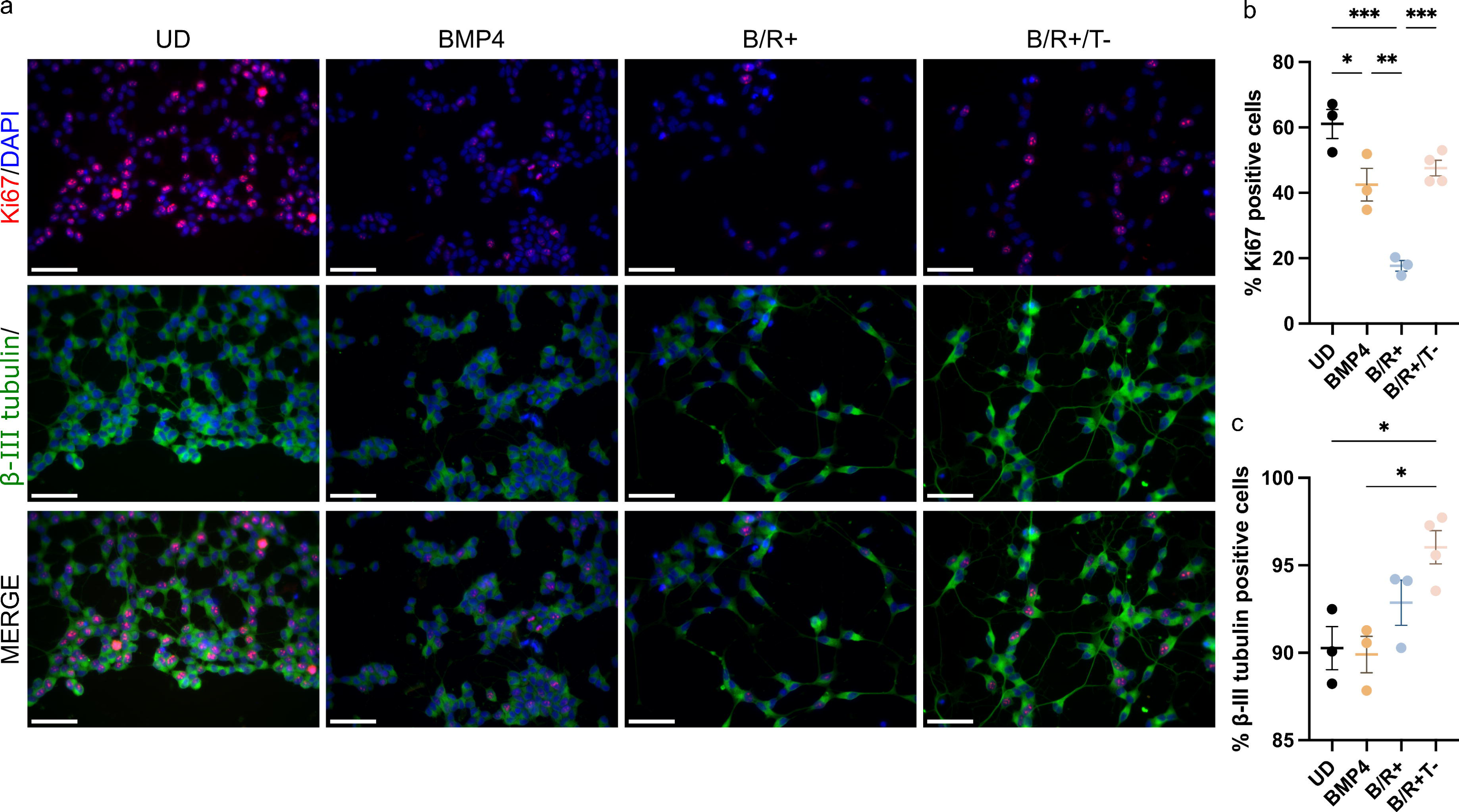
The BRT differentiation of SH-SY5Y cells leads to significant changes in the levels of β-III tubulin and Ki67 positive cells. a. DAPI, Ki67 and β-III tubulin staining of SH-SY5Y cells at different time point along the B/R+/T− induced differentiation. b-c. Quantification of the percentage of Ki67 (b) and β-III tubulin (c) positive cells for B/R+/T− treated SH-SY5Y cells at different time points along the differentiation process. One-way ANOVA to compare means of UD, BMP4, B/R+ and B/R+/T−, *p<0.033, **p<0.002, ***p<0.001 (see Table S1 for p-values and statistical analysis details). Data shown as mean ± SEM and derive from the analysis of 3 (UD, BMP4, B/R+) or 4 (B/R+/T−) images. Scale bar = 50 µm.

In the light of our shotgun lipidomic, neurite length, ICC and WB analyses, we investigated potential correlations among the levels of lipids significantly affected by the differentiation process (Figure 1 and 2 and Table 1) and TH or neurite length (Figure 3b and d). We found that the levels of DAG, SM, Cer and HexCer correlate positively whereas those of PS correlate negatively with those of TH and/or neurite length (Figure 5a-d). Moreover, we found that the levels of the SL species that were the most systematically affected by the differentiation all correlate positively (36:1) or negatively (40:1) with those of TH and/or neurite length (Figure 5e-n). Finally, we found that the levels of all most abundant DAG and glycerophospholipid species (levels higher than 5%, shown in bold in Table 1), except PCO-34:2, PA 44:5, correlate with those of TH and/or neurite length (Figure S14).

**Figure 5:**
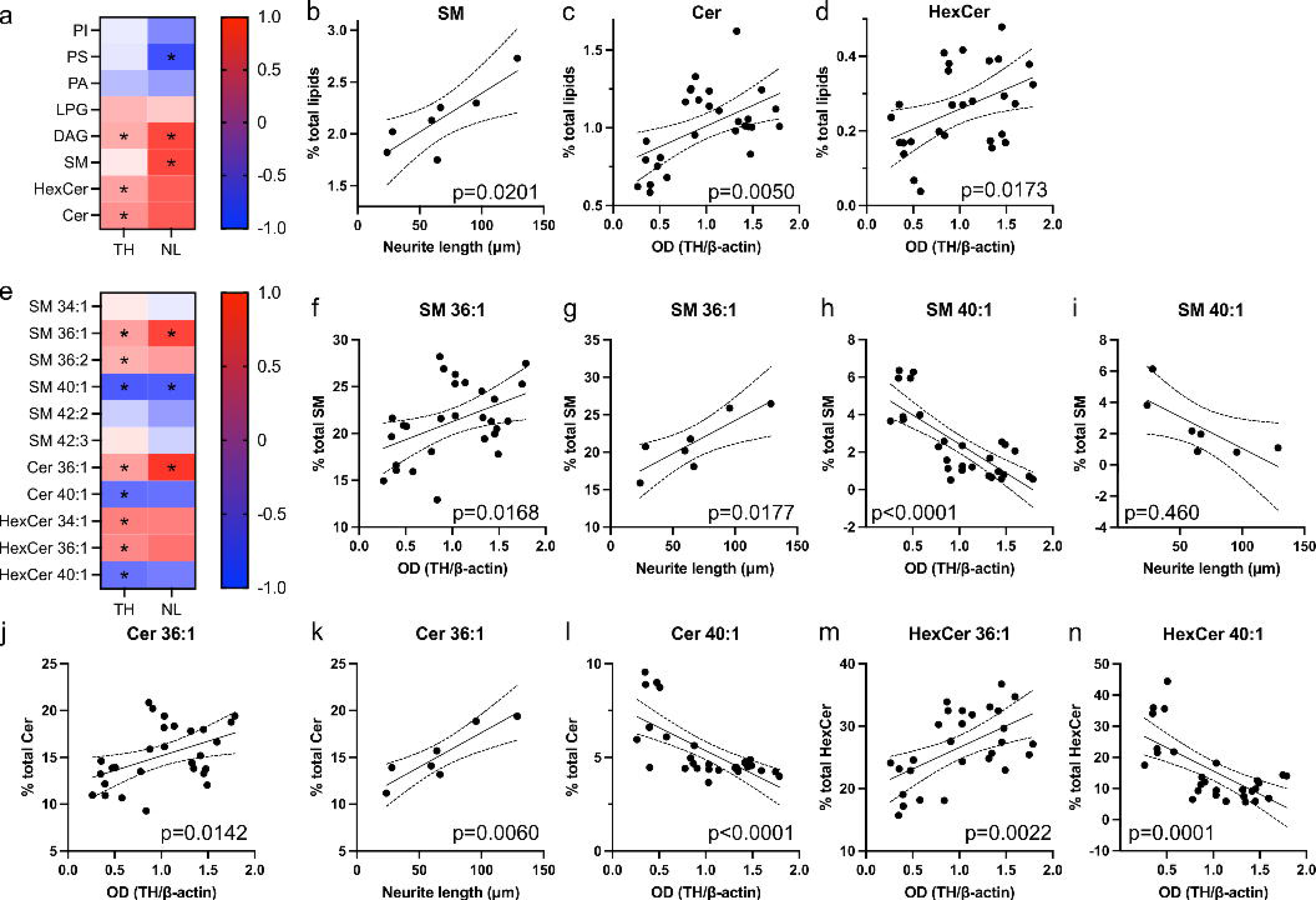
The levels of sphingolipids and their 36:1 and 40:1 subspecies correlate with those of TH and/or neurite length. a,e. Heat maps representing Pearson r for the correlation between the lipid class (a) and percentage of phospholipid subspecies (e) vs OD (TH/β-actin) (TH) or neurite length (NL) for the data sets UD, BMP4, B/R+, B/R-, B/R+/T+, B/R+/T−, B/R−/T− (*p<0.033). The main subspecies for each lipid class are shown in bold. b-d, f-n. Linear regressions showing the correlation between lipid classes or lipid subspecies and TH (28 data points) and/or neurite length (7 data points).

Altogether, our combined lipidomic, neurite length, ICC and WB analyses, show that the levels of lipid species significantly affected by the differentiation of SH-SY5Y cells correlate with those of neuronal and/or dopaminergic markers.

### BRT-induced differentiation affects the levels of proteins involved in lipid metabolism and processes associated with neuronal maturation

We further analysed the proteome changes of undifferentiated and BRT-differentiated SH-SY5Y cells to investigate potential changes in proteome associated with the differentiation process by mass spectrometry (MS, Figure 6 and S15). We could identify 149 differentially expressed proteins between B/R+/T− differentiated SH-SY5Y cells compared to undifferentiated cells (Figure S15b). Our proteomic data show that the BRT-induced differentiation led to significant changes in the levels of proteins involved in biological processes (GO) associated with neuron maturation, such as cell adhesion, locomotion, cell motility, cell migration, and lipid metabolic processes (Figure 5a and S15) and thus strongly support our lipidomic, WB, neurite length and ICC analysis. In particular, the increase in neurite length associated with the BRT-induced differentiation reported in Figure 3 can be supported with the observed increase in the levels of Microtubule-associated protein tau (MAPT), a protein mainly found in axons in mature neurons under physiological conditions^62,63^, cell adhesion molecule 3 (CADM3), that belongs to the CADM protein family known to be involved in neural network formation, such as axon-guidance, synapse formation and regulation of synaptic structure^64^, Galactin 1 (LGALS1), a protein involved in cell-cell interactions and dihydropyrimidinase like 3 (DPYSL3), that belong to the DPYSL protein family involved in neurite extension^65^, axonal guidance^66,67^ and synaptic plasticity^68^. Our proteomic data also show that BRT-induced differentiation led to a tendency towards increased levels of β-III tubulin and a significant increase in those of αS which is in agreement with our ICC analyses but in contrast to our WB analyses. The discrepancy between the results of the proteomics and WB results may come from the high variability associated with the WB analyses. Finally, our proteomic data confirmed the absence of changes in the levels of Ki67 seen in our ICC analysis upon BRT-differentiation of SH-SY5Y cells but showed the decrease in the levels of other proteins involved in cell proliferation such as Protein kinase C alpha (PRKCA)^69,70^, as expected and previously reported for differentiated cells^71^.

**Figure 6:**
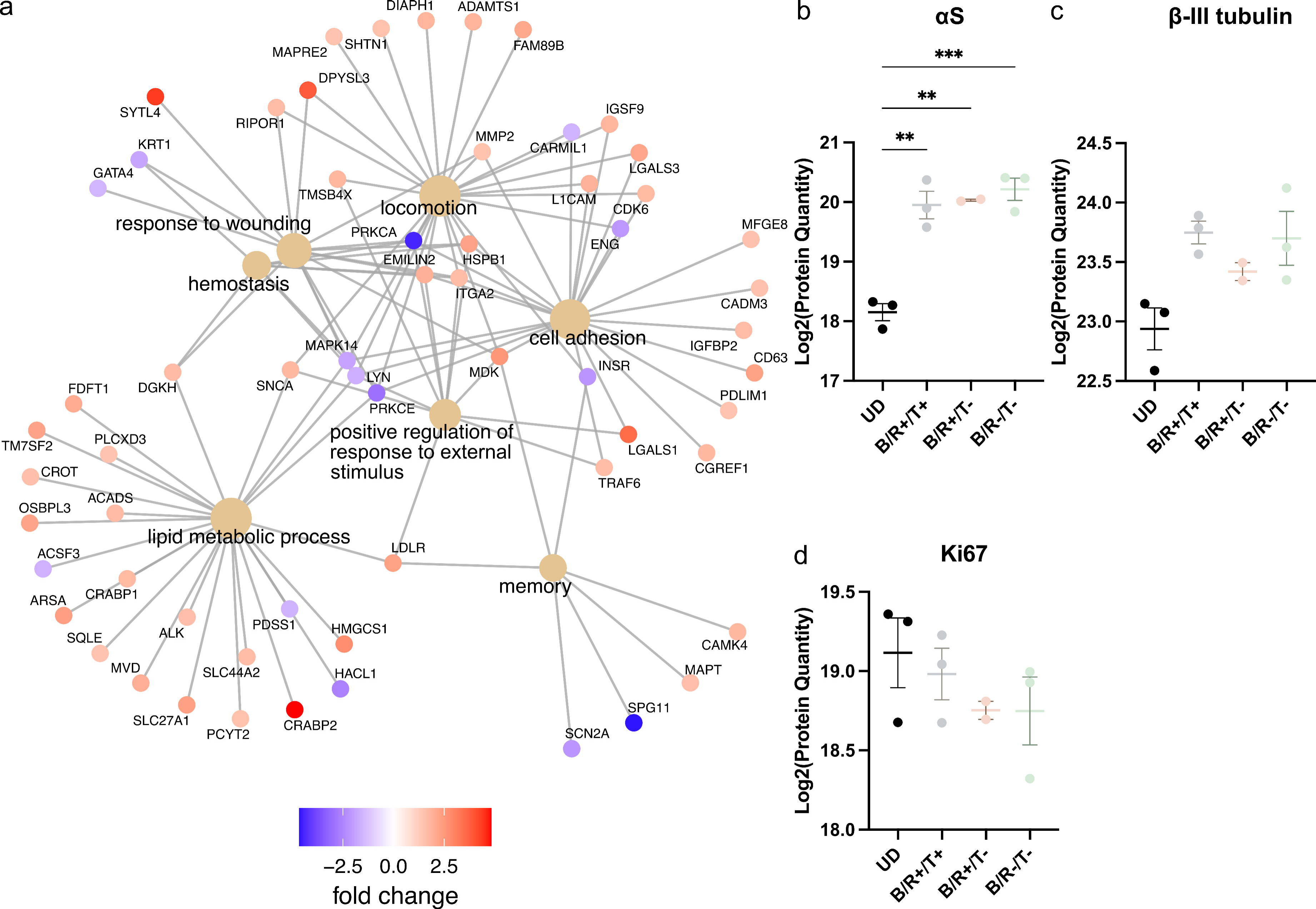
BRT-induced differentiation of SH-SY5Y cells leads to changes in the levels of proteins involved in lipid metabolism and neuron maturation. (a) Biological function GO over-representation analysis of the differentially expressed proteins between undifferentiated and BRT-induced differentiated SH-SY5Y cells. The 15 most overrepresented GO term are displayed with their differentially expressed proteins. Direct children of overarching GO terms are excluded. All 15 GO-terms shown in Figure S15 (b-d) Log2 protein quantities for αS, β-III tubulin and Ki67 in undifferentiated and BRT-induced differentiated SH-SY5Y cells. Mean highlighted as dash. Significant differences based on adjusted p-values of t-test (* p<0.05, ** p <0.01, *** p<0.0001). Data shown as mean ± SEM and derive from 2 (B/R+/T−) or 3 (UD, B/R+/T+, B/R−/T−) independent differentiations.

### BR and BRT treatments are required for the cells to mature to a sufficient neuronal phenotype capable of eliciting action potentials

Finally, we measured electrophysiological features of SH-SY5Y cells along the differentiation process by monitoring resting membrane potential, current necessary to hold the cells at -70 mV, rheobase, action potential (AP) amplitude, and the amplitude of the AP after hyperpolarisation (AHP) in SH-SY5Y cells at different time points along the differentiation process (Figure 7). We measured a resting membrane potential for the undifferentiated cells of ca. - 26mV, a value similar to those previously reported^72^. No significant changes in the resting membrane potential and holding current required to hold the cells at -70 mV was observed between undifferentiated and differentiated cells (Figure 7a,b). We could elicit APs (Figure 7c1) in BR and BRT differentiated cells but not in undifferentiated and BMP4 treated SH-SY5Y cells, even at very depolarised potentials (Figure 7c2). However, we did observe spikelet type APs in all treatment groups and in the undifferentiated cells, which were categorized as no APs (Figure 7c3). There were no differences in the proportion of cells in which APs could be elicited when comparing B/R+ and B/R-, B/R+ with B/R+/T+ and B/R+/T− or B/R- with B/R−/T−. No differences were observed in rheobase as the current required to elicit an AP did not significantly differ between the groups (Figure 7d). We however observed that the final TPA treatment led to a significant increase in the AP amplitude in the presence of serum and significant increase in AHP amplitude when comparing B/R+/T+ vs B/R+ treated cells (Figure 7e,f).

**Figure 7:**
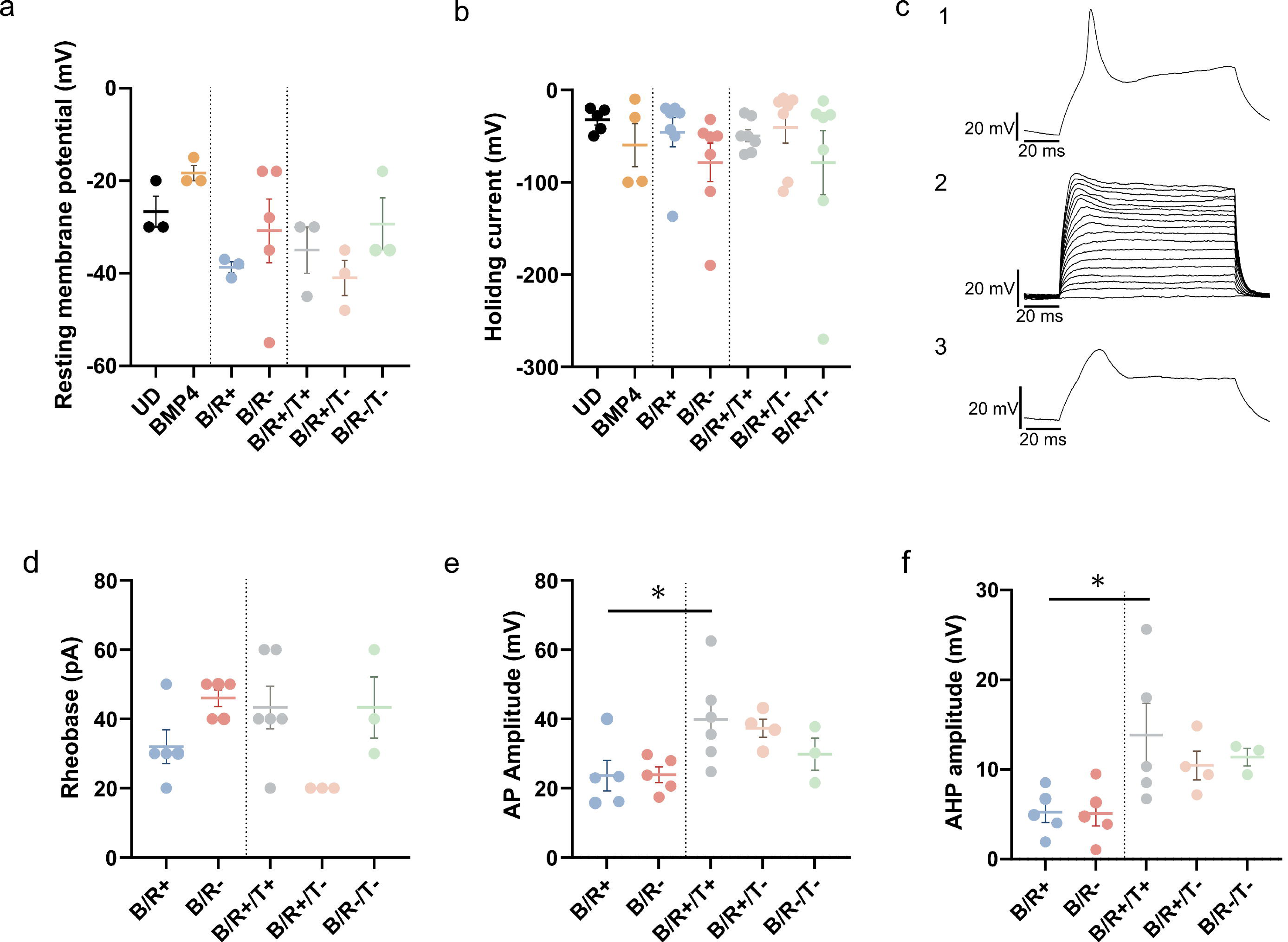
The BRT differentiation of SH-SY5Y cells promotes electrical properties. a-b. Scatter plots showing the resting membrane potential (a) and holding currents (b) of undifferentiated cells and BMP4, B/R+, B/R-, B/R+/T+, B/R+/T− and B/R−/T− treated cells. c. Representative membrane responses of a differentiated cell in which an AP could be elicited in response to injection of positive current in current clamp mode (c1), injection of incrementing positive current steps (10 pA) in current clamp mode failed to elicit APs in undifferentiated- and differentiated cells (c2); spikelets could be elicited in both cell types (c3). d-f. Rheobase (d), amplitude of the APs (e) and amplitude of the after hyperpolarization (f) measured for B/R+, B/R-, B/R+/T+, B/R+/T− and B/R+/T− treated cells are plotted. One-way ANOVA to compare means of UD, BMP4, B/R+, B/R-, B/R+/T+, B/R+/T− and B/R−/T− (a-b), or means of B/R+, B/R-, B/R+/T+, B/R+/T− and B/R−/T−. *p<0.033. Comparison shown: BMP4, B/R+/T+, B/R+/T− and B/R-T- vs UD, B/R+ and B/R- vs BMP4, B/R+/T+, B/R+/T− and B/R−/T− vs their respective B/R(+/-). (e: One-way ANOVA: p = 0.0310; f: (One-way ANOVA: p = 0.0236)).

Altogether, these results show that BR and BRT treatments are required for the cells to mature to a sufficient neuronal phenotype to show AP elicitation. Moreover, our combined shotgun lipidomic, neurite length, WB and AP measurements show that the final TPA treatment is required for lipid and TH levels as well as neurite length to reach their maximal/minimal values and for the detection/measurement of an AP and AHP amplitude in differentiated SH-SY5Y cells. These results suggest the potential requirement of a minimal concentration of specific lipids in the membranes of differentiated SH-SY5Y cells for the detection of an electrophysiological activity in these cells.

## Discussion

PD is a neurodegenerative disease characterised by the death of dopaminergic neurons in the substantia nigra pars compacta and the deposition of αS-containing protein aggregates in the remaining cells^1^. SH-SY5Y cells is the most commonly used *in vitro* model to study PD and αS pathology as well as the environmental and genetic risk factors of the disease due to the fact that these cells express dopaminergic markers and can be differentiated into more neuronal and dopaminergic-like cells^24–38^. Undifferentiated SH-SY5Y cells are proposed to resemble immature catecholaminergic neurons^43^ and express cell proliferation and immature neuronal markers^47,54^. Differentiation of SH-SY5Y cells using subsequent incubations with RA and TPA was found to lead to a more mature dopaminergic-like phenotype thus making RA/TPA-differentiated SH-SY5Y cells a robust model for PD^28,32,48,59^. Although increasing evidence points towards a role of lipids in the initiation and development of the disease, limited data are currently available on the lipidome of these cells.

The results of this study provide a unique quantitative and comprehensive description of the changes in the lipidome of SH-SY5Y neuroblastoma cells associated with an increase in neuronal and dopaminergic markers as well as electrophysiological activity along the differentiation process. We induced the differentiation of SH-SY5Y cells using subsequent treatments with BMP4, RA and TPA in the presence (+) or absence (-) of serum (B/R+/T+, B/R+/T− and B/R−/T−). BMP4 pre-treatment was used to enrich the N-type subpopulation at the expense of the S-type^42^. Our combined brightfield and fluorescence imaging as well as WB analyses show that undifferentiated SH-SY5Y cells express relatively low levels of αS, TH and D3R, have short neurites (ca. 25 μm) and are positive to both Ki67 (60%) and β-III tubulin (90%) staining. These observations are in agreement with previous studies that have reported that undifferentiated SH-SY5Y express dopaminergic, cell proliferation and immature neuronal markers^47,54^. We observed that BMP4 treatment decreased the proliferation of the cells, led to a more homogeneous cell population and affected the levels of SL species to a limited extent but did not affect the neurite length, TH and D3R levels or the percentage of β-III tubulin cells (Figure 8). Both undifferentiated and BMP4 treated SH-SY5Y cells were found to have resting membrane potential of ca. -25 mV and not to elicit action potential, as reported previously^73^. Further treatment of SH-SY5Y cells with RA did not affect the levels of D3R but led to a significant increase in the levels of αS and TH as well as neurite length and a significant decrease in the proliferation of the cells, in agreement with previous studies that used RA treatment in combination with low FBS to induce the differentiation of SH-SY5Y cells^47^ (Figure 8). These BR-induced changes in dopaminergic, neuronal and proliferation markers were associated with significant changes in the levels of some lipid species, e.g. 40:1 SL and most of DAG species as well as the ability of the cells to elicit AP (Figure 8). Only the BRT treatments that consist in the subsequent incubations of SH- SY5Y cells with BMP4, then RA then TPA were found to lead to the significant increase of all studied neuronal (neurite length, β-III tubulin positive cells) and dopaminergic (TH and D3R) markers as well as the maximal observed change in the levels of lipids affected by the differentiation process (Figure 8). Proteomic analyses confirmed that the BRT-induced differentiation of SH-SY5Y cells significantly affect lipid metabolism and neuron maturation processes. In particular, we reported increased levels of neuronal markers (e.g. MAPT) and a decrease in the levels of proteins involved in cell proliferation (e.g. PRKCA) in BRT-differentiated SH-SY5Y cells compared to undifferentiated cells, as reported previously for RA-, RA/BDNF- or RA/TPA-differentiated cells^49,71,74^. Finally, we found that both BMP4 pre-treatment and the removal of serum during the RA and/or TPA led to a more pronounced increase in neurite length, as described previously for serum removal^75^, and more pronounced changes in the levels of lipid species associated with SH-SY5Y cells differentiation.

**Figure 8:**
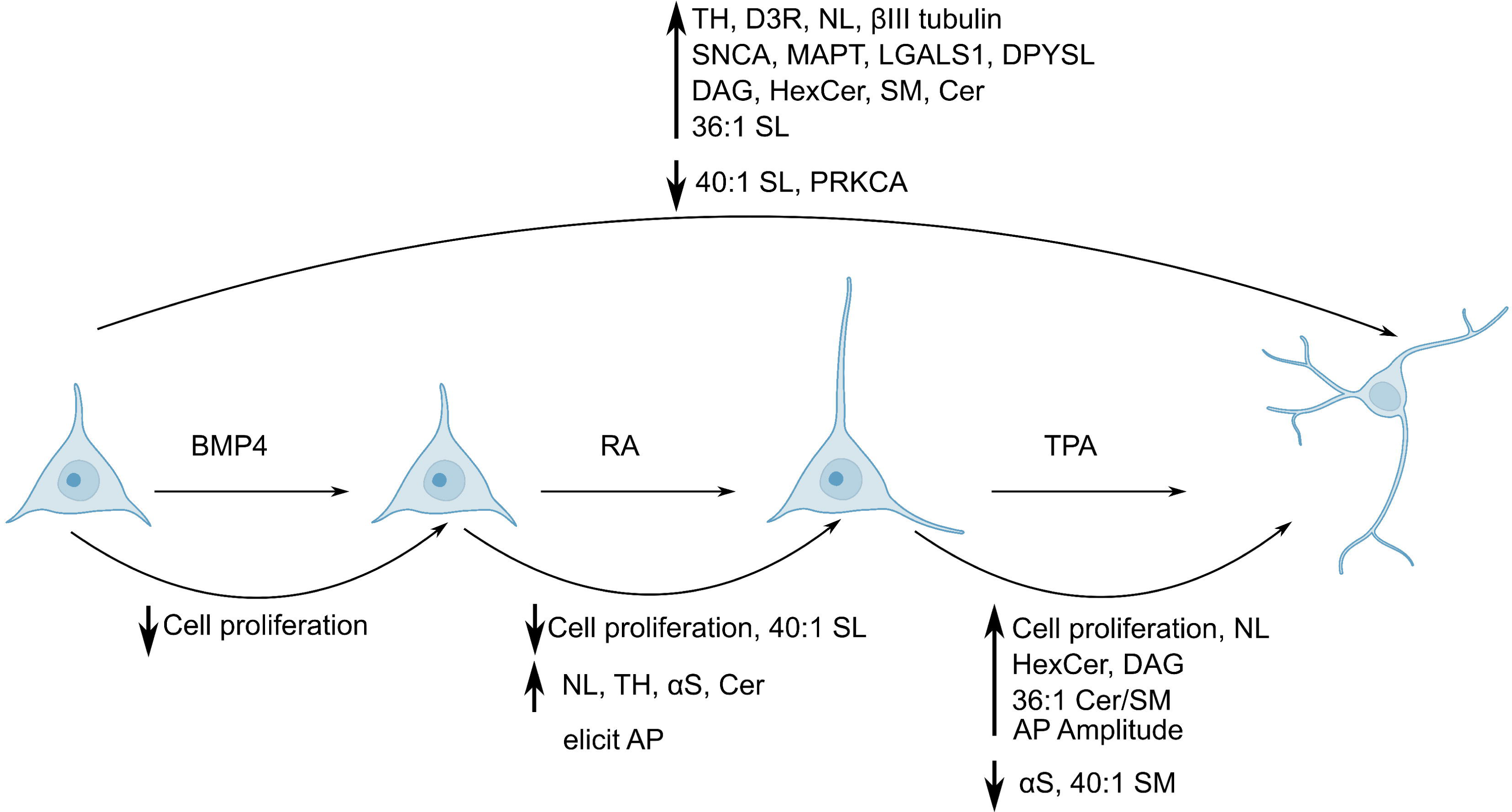
Overview of the phenotypic changes induced by the BRT differentiation. Changes associated with the complete treatment and after each step of the treatment are shown above and below the illustration, respectively. Figure prepared with Biorender.

Our combined shotgun lipidomic, biochemical and microscopic analyses show that BRT-induced differentiation of SH-SY5Y cells was associated with significant changes in the levels and chain distribution of DAG, PS, Cer, SM and HexCer and that their levels and those of their species correlate with those of TH and/or neurite length. SL were the class of lipids most affected by the differentiation process in term of their levels and chain properties as expected given the fact that SL are a class of membrane lipids that are involved in signalling, cell differentiation, proliferation and apoptosis^76,77^. The BRT-induced differentiations also led to an increase in the levels of the 36:1 species and the decrease in those of the 40:1 species for HexCer, SM and Cer, as the cells became more neuronal and dopaminergic-like (Figure 8). The chain distribution of SL was found to be specific to the type of brain cells^78^ (*e.g.* astrocyte, microglia, neurons and oligodendrocytes) with neuronal cells being particularly enriched in 36:1 SL species and poor in 40:1 SL species (Figure S16a-c, data from Fitzner et al^78^). Moreover, the maturation of primary neurons for 5, 10 and 16 days *in vitro* was found to be associated with an increase in the 36:1 SM and Cer species (Figure S16d-e, data from Fitzner et al^78^), suggesting that the increase in the level of this particular SL species may contribute to a lipid signature of neuron maturation. Even though the percentage of 36:1 SL in the BRT-differentiated SH-SY5Y neuroblastoma cells are lower than that observed in primary neurons (20-40% in BRT- differentiated SH-SY5Y cells vs 60-80% in primary neurons after 15 days *in vitro*), our observation that the levels of the 36:1 SL species increase and those of the 40:1 SL species decrease as the cells become more neuronal is in line with the reported chain distribution of SL in neurons and the increase in 36:1 SL species associated with their maturation *in vitro*^78^. In addition to the biological relevance of SL in the context of cell differentiation, these lipids are also relevant to PD as their chains were found to be altered in association with both the sporadic and familial form of the disease. Indeed, lipidomic analyses of post-mortem brain samples and fibroblasts of PD patients have shown the accumulation of short chain species for Cer, SM and HexCer in patients’ samples ^18,19^. Given the fact that changes in the levels and chain properties of SL have been reported in association with PD, there is a strong need for a thorough characterisation of the lipidome of the cell model used for the study of PD as well as the lipidome changes associated with the differentiation process in order to to discriminate lipid changes associated with PD and those associated with the differentiation of the cells into more mature and dopaminergic-like phenotype.

In conclusion, we have provided the quantitative description of lipid changes associated with the differentiation of SH-SY5Y neuroblastoma cells into more mature and dopaminergic-like cells. The levels of the lipids affected by the differentiation process were found to correlate with the levels of TH and/or neurite length. The observed changes in lipidome and neuronal markers associated with the differentiation of SH-SY5Y cells are supported by proteomic analyses showing that the differentiation mainly affected biological processes associated with lipid metabolism and neuronal maturation. Our study provides the complete and quantitative characterisation of the changes in lipidome associated with the differentiation of SH-SY5Y cells into more neuronal and dopaminergic-like phenotype and serves as a basis for further characterisation of lipid disruptions in association with PD and its risk factors in this dopaminergic neuronal cell model.

## Materials and methods

### Cell culture

SH-SY5Y cell line (ATCC, CRL-2266TM) was maintained according to recommended guidelines in a humidified incubator at 37°C and 5% CO2 and tested negative on mycoplasma. Cells were maintained in Dulbecco’s modified eagle medium (DMEM), high glucose, Glutamax (Thermofisher Scientific, 31966021) supplemented with 1% v/v non-essential amino acids (NEAA, Thermofisher Scientific, 11140050), 1% v/v Penicillin-Streptomycin (P/S, Thermofisher Scientific, 15140122) and 10% v/v Fetal Bovine Serum (FBS) (Sigma-Aldrich, F7524). Medium was changed every three days and cells passaged and re-cultured when 80% confluent. Pretreatment with BMP4 was carried in maintenance media with 5% v/v FBS, supplemented with 100 ng/ml BMP4 (Miltenyi Biotec, 130- 111-168) and media was changed every three days. Differentiation of SH-SY5Y cells was carried out at a cell density between 17000-35000 cells/cm2 in Nunclon™ Delta surface treated polystyrene flasks or dishes (Thermofisher Scientific) or 35 mm Petri dishes for elecrophysiology. Differentiation medium was 1:1 v/v DMEM/F-12 (Thermofisher Scientific, 21331020) and NaurobasalTM medium (Thermofisher Scientific, 21103049) supplemented with 0.5% v/v L-Glutamine (Thermofisher Scientific, 25030032), 0.5% v/v N-2 supplement, 1% v/v FBS and 1% v/v P/S. Additional supplementation of medium during differentiation with retinoic acid (RA, Sigma-Aldrich, R2625) or 12-O-tetradecanoylphorbol-13-acetat (TPA, Sigma-Aldrich, P8139) was done at 10 µM and 80 nM, respectively, with media change every three days.

### Neurite length measurement

Cell culture images were taken every 24 h on Zoe Fluorescent cell imager (Bio-Rad). Three individual images were taken per condition and timepoint. Neurite length for each image was measured via previously described protocol using the NeuronJ application in ImageJ Software (Pemberton et. al., 2018). A neurite was defined as any projection protruding from a cell body (soma).

### Western blot

Cells were washed with ice-cold phosphate-buffered saline (PBS) and lysed in extraction buffer (1% (v/v) triton X-100 and protease inhibitor (Sigma-Aldrich, 11836170001) in PBS). Cell lysates were incubated on ice for 10 min and centrifuged at for 10 min at 20,000 g and 4°C. Supernatant was collected and protein content was measured using the PierceTM BCA Protein Assay Kit (Thermofisher Scientific, 23225).

Lysates containing 25 µg of protein were electrophoresed on a NuPage 4%-12% Bis-Tris Protein gel (Thermofisher Scientific). Proteins were transferred to a PVDF membrane (0.2 µm, Thermofisher Scientific, 88520), blocked in 5% w/v milk (2 h) and treated with primary and secondary antibodies. Antibody binding was detected using an ECL chemiluminescence kit (Bio-Rad, 1705061). The following antibodies were used: α-synuclein (BD Biosciences, 610787, 1:500), β-actin (ABclonal, AC026, 1:100.000), dopamine D3 receptor (Merck, AB1785P, 1:1000), TH (Cell Signaling Technology, #2792, 1:1000), anti-mouse (Abcam, ab205719, 1:5000) and anti-rabbit (Abcam, ab6721, 1:5000).

### Immunocytochemistry measurements

Cells used for immunocytochemistry were plated on 48-well plates coated with laminin 521 (1 µg/cm2). SH-SY5Y cells were fixed at different time points of the differentiation by incubating cells with 4% paraformaldehyde (15 min, 22°C) followed by three washes with PBS. Cells were permeabilized with blocking buffer (0.2% Triton-X and 5% Donkey Serum in PBS, 1.5 h, 4 °C). Cells were then incubated with primary antibodies (ON, 4 °C, in blocking buffer) targeting Ki67 (abcam, ab16667, 1:250) and β-III-tubulin (abcam, ab78078, 1:1000), followed by three washes with PBS. Finally, cells were incubated with secondary antibodies (2 h, 22°C, protected from light) which were either anti-rabbit Alexa Fluor 594 (Jackson Immunoresearch, #711-585-152, 1:200) and anti-mouse Alexa Fluor 488 (Jackson Immunoresearch, #715-545-150, 1:200) mixed with DAPI (Sigma, MBD0015, 1:500) followed by three washes with PBS. At least three pictures were taken from each well using a Leica DM IL LED Fluor Inverted epifluorescence microscope and analysis was performed using LASX software and CellProfiler^79^.

### Lipid extraction and shotgun lipidomics

Cells were washed with ice-cold PBS, spun down (5 min, 4°C, 2000 g), and pellet was washed once with PBS and another time with 1 ml ice-cold 155 mM ammoinium bicarbonate (AB, Sigma-Aldrich, FL40867) until finally resuspended in same buffer to 5000 cells/µl. Lipid extraction and shotgun lipidomics was done using quantitative MS-based shotgun lipidomics as previously described^80^ with minor modifications. Full lipid extraction was performed on a fraction containing approximately 3*10^5^ cells per sample. Each cell suspension was spiked with a 10x Internal Standard Mix (IS mix) with a known content of a variety of different lipids^80^. Organic solvent containing mixture of chloroform and methanol (C/M, 2:1 v/v, 1 ml) was added to the cell suspension and suspension was mixed (15 min, 4°C, 2000 RPM) and centrifuged (2 min, 4°C, 2000 g). The organic phase was isolated, methanol (100 µl) and AB (50 µl) added and solution was once again mixed (5 min, 4°C, 2000 RPM) and centrifuged (2 min, 4°C, 2000 g). Organic phase was transferred into a new tube and evaporated for 1 h. The lipid film was dissolved in C/M (1:2 v/v, 100 µl) by mixing (5 min, 4°C, 2000 RPM) and then centrifuged (15 min, 4°C, 15000 g). Each lipid extract was mixed with positive or negative ionization solvents (13.3 mM ammonium bicarbonate in 2-propanol or 0.2% (v/v) tri-ethyl-amine in chloro-form/methanol 1:5 (v/v), respectively) and analyzed in the positive and negative ion modes on quadrupole-Orbitrap mass spectrometer Q Exactive (Thermofisher Scientific) equipped with TriVersa NanoMate (Advion Biosciences) for automated and direct nanoelectrospray infusion.

The data acquisition cycle consisted of FT MS and FT MS/MS scans in the positive and negative ion modes. The machine settings were as described previously^81^. Initial processing of the acquired FT MS and FT MS/MS data that reported identified lipid species together with values of m/z and intensities for the associated precursor and fragment ions was performed using the LipidXplorer software^82^. Lipids were identified (to Metabolomics Standards Initiative level 1^83^ when the measured m/z values of precursor ions and associated fragment ions fulfilled the criteria defined in molecular fragmentation query language (MFQLs) for known lipid classes, as listed in Table S6 in Supplementary Materials. The obtained list of identified lipid species were examined manually to remove those improperly quantified e.g. due to noise. An in-house build R-based suite of scripts named LipidQ (https://github.com/ELELAB/lipidQ) was used for the calculation of the absolute molar quantities of the identified lipid species in samples based on the reported intensities of sample-derived lipids and internal lipid standards.

### Whole-cell patch clamp electrophysiology

SH-SY5Y cell media was replaced on the day of recordings with artificial balanced salt solution (ABSS) at room temperature (20-24 °C). The ABSS solution consisted of (in mM): 140 NaCl, 3.5 KCl, 1.25 Na2PO4, 2 MgSO4, 2 CaCl2, 10 Glucose, and 20 HEPES at pH: 7.35. The dish was placed on an Axiovert 10 microscope (Zeiss) under constant perfusion with fresh ABSS media, and cells were observed through x200 magnification. Whole-cell patch clamp recordings of the cells in voltage- and current mode were made using an EPC-9 amplifier (HEKA), and micropipettes (1.5 mm OD Glass) (World Precision Instruments) fabricated with a microelectrode puller Model PP-830 (Narishige). Pipettes exhibited a 2.0 - 4.0 MΩ resistance when filled with an intracellular solution which contained (in mM, pH of 7.3): 130 K-Gluconate, 10 KCl, 10 Hepes acid, 1 MgCl2, 1 CaCl2, 10 EGTA and 2 MgATP was used to measure the resting membrane potential, or 140 KCl, 1 MgCl2, 1 CaCl2, 10 EGTA, 2 MgATP, and 10 HEPES to evaluate holding currents, rheobase and action potential (AP) parameters. Cells were held at -70 mV and a 70 % compensation of the series resistance was applied. Current and voltage relationships were obtained by holding cells at -70 mV for 20 ms before and after 200-300 ms of incrementing voltage steps starting at -110 mV with an increase of 10 mV until the stepped potential reached +20 mV. Current and voltage relationships along with AP elicitation made in current clamp were obtained by applying incrementing steps of injected current (+10 pA steps; 25-30 steps) in the cell from the amount of current necessary to hold the membrane potential of the cell at -70 mv. In some cases, while APs could not be elicited, a small, spike like depolarization with a small or absent afterhyperpolarization (AHP), could be elicited which was designated as a spikelet and not included as an AP in analysis. Recordings were analyzed in Pulse and PulseFit software 8.8 (HEKA) and figures were made using Igor Pro software 6.20 (Wavemetric) and Graphpad Prism™ 9.0.

A 2x2 contingency table was used when calculating statistical differences in the proportion of cells eliciting an AP, while differences in resting membrane potential, holding currents and AP parameters were analysed using a one-way ANOVA test using Graphpad.

### Proteomics

Cells were washed twice with ice-cold PBS, spun down (8 min, RT, 4000 g), and lysed in 6M Guanidinium Hydrochloride, 50mM HEPES pH 8.5, 40mM 2-chloroacetamide and 10mM Tris(2- carboxyethyl)phosphine, boiled at 95 °C for 5mins and sonicated for 3x cycles at 30 sec on, 30 sec off in a Bioruptor Pico sonicator. Before digestion, the lysate was diluted 10x with 50 mM HEPES, 10 % Acetonitrile and digested with Trypsin/LysC (Thermo Scientific™) in a 1:50 (enzyme-to-protein) ratio. Cell lysate was digested over night at 37 °C and quenched with TFA to a final concentration of 0.5 %. Digest was desalted using a Thermo Scientific™ SOLAµ™ Solid Phase Extraction (SPE) plate and the resulting eluate was vacuum dried at 45 °C. For each sample, 500 ng digest was loaded on Evotips (EvoTip Pure) and analyzed on a Thermo Fisher Exploris 480 mass spectrometer combined with FAIMSPro and an EvoSep One chromatography system. Samples were acquired with a 20 samples-per-day (SPD) Whisper method (58 min) in randomized format with an Aurora Elite TS Generation 3 15 cm column (IonOptics). The samples were acquired in DIA with the FAIMS Pro interface set to a compensation voltage of -45V. MS1 scans were performed at 120k resolution with an AGC of 300 % and an injection time of max 246 ms. Precursors were fragmented in HCD with a NCE of 32 %. MS2 scans were acquired at 120k and over 59 windows, each spanning 10 m/z across 400-1000 m/z. An MS1 scans were inserted every 20 MS2 scans.

Raw files were searched with Spectronaut 17. Trypsin and LysC were selected as protease and quantification was performed at MS1 level. All other parameters were kept at default. Further data analysis was performed in R. Only proteotypic and protein-group-specific peptides were included and peptides with and MS1 or MS2 quantity below 50 were excluded. Precursors with charge state <2 were excluded. Protein quantities for the remaining peptides were extracted. One replicate of B/R+/T− was excluded due to irreproducible digestion. Protein identifications in single replicates were removed. No protein quantities were imputed and only proteins identified in all replicates were considered for PCA and differential expression analysis. Differential expression analysis was performed with the limma package and p-values were adjusted by Benjamini-Hochberg^84,85^. Significance thresholds were set to 0.05 for the adjusted p-value and 1.5 for the log2(Fold-change). Differentially expressed proteins were further passed on to clusterProfiler for an overrepresentation analysis based on the Gene Ontology (GO) terms biological processes (BP)^86,87^. The p-value was adjusted by Benjamini-Hochberg with a cutoff of 0.01. Output was visualized using the enrichplot package. The 15 most overrepresented GO terms were illustrated and further manually curated by excluding child terms of higher present GO terms. All GO terms can be found in the supplementary material.

### Statistical analysis

Statistics were performed, and graphs were generated, using Prism 9 software (GraphPad). Statistical significance was determined using One-way ANOVA and multiple t-test comparisons (Šídák’s multiple comparisons test). Data were shown as means ± SEM for all results. Effects achieving p < 0.033 were interpreted as statistically significant. No samples were excluded from any analysis unless explicitly stated. Comparisons were performed to assess the effect of the BRT- induced differentiation (UD vs B/R+/T+ or B/R+/T− or B/R−/T−) and of the individual treatment, i.e. BMP4 (UD vs BMP4), RA (BMP4 vs B/R- or B/R+), TPA (BR- vs BR-/T− or B/R+ vs B/R+/T+ or B/R+/T−). The details of the statistical analyses for all figures are shown in Table S1.

## Acknowledgements

This work was supported by the Independent Research Fund Denmark, Sino-Danish Center for Education and Research, Lundbeck Foundation, Carlsberg Foundation, Novo Nordisk Foundation and Parkinsonforeningen.

