## Supplementary Information for "Lipid levels correlate with neuronal and dopaminergic markers during the differentiation of SH-SY5Y cells"

### Supplementary Figures

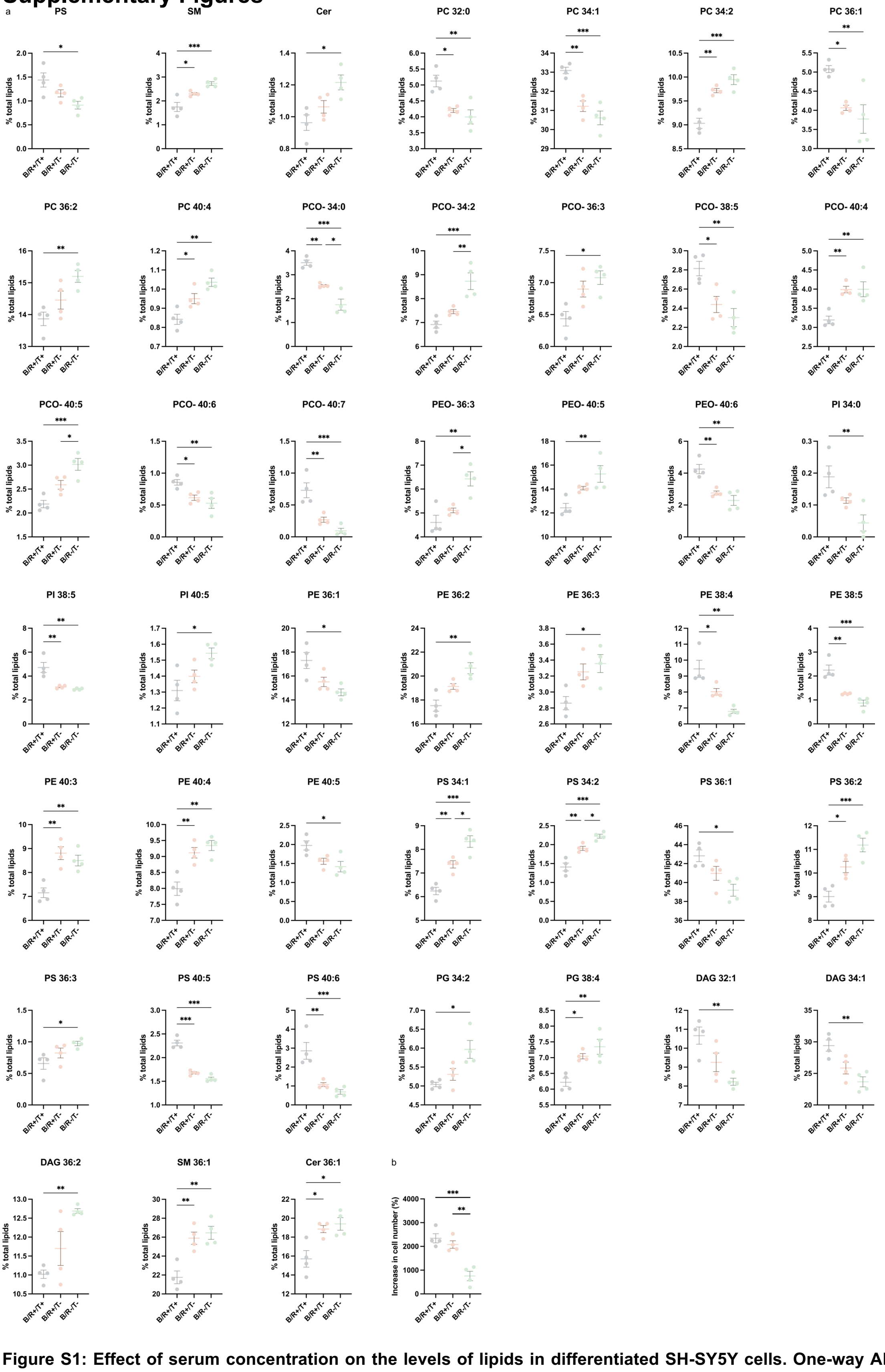


**Figure S1: Effect of serum concentration on the levels of lipids in differentiated SH-SY5Y cells. One-way ANOVA to compare means of B/R+/T+, B/R+/T- and B/R-/T-, *p<0.033, **p<0.002, ***p<0.001 (see Table S1 for p-values and statistical analysis details). Data shown as mean ± SEM and derive from 4 independent differentiations.**

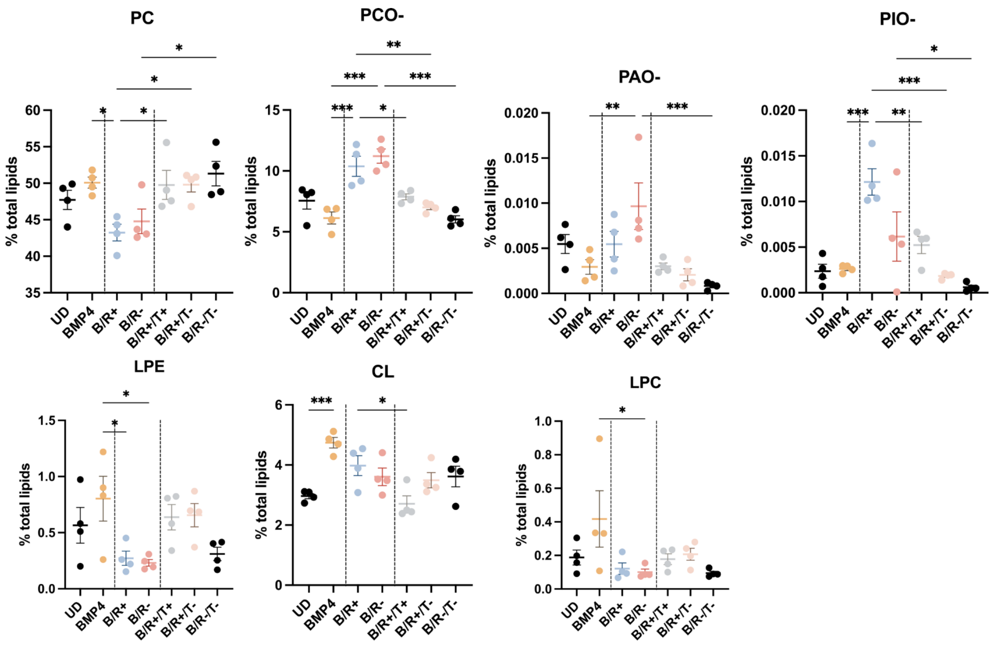


**Figure S2: Lipids whose levels are affected significantly at the BMP4 or RA stage but do not differ when comparing UD and BRT cells. Only species whose levels are significantly affected at the BMP4 or RA stage and that have been detected in all UD and BRT replicates are shown. One-way ANOVA to compare means of UD, BMP4, B/R+, B/R-, B/R+/T+, B/R+/T- and B/R-/T-, *p<0.033, **p<0.002, ***p<0.001. Comparisons shown: UD vs all, BMP4 vs B/R+, BMP4 vs B/R-, B/R+ vs B/R+/T+, B/R+ vs B/R+/T-, B/R- vs B/R-/T-. Data shown as mean ± SEM and derive from 4 independent differentiations.**

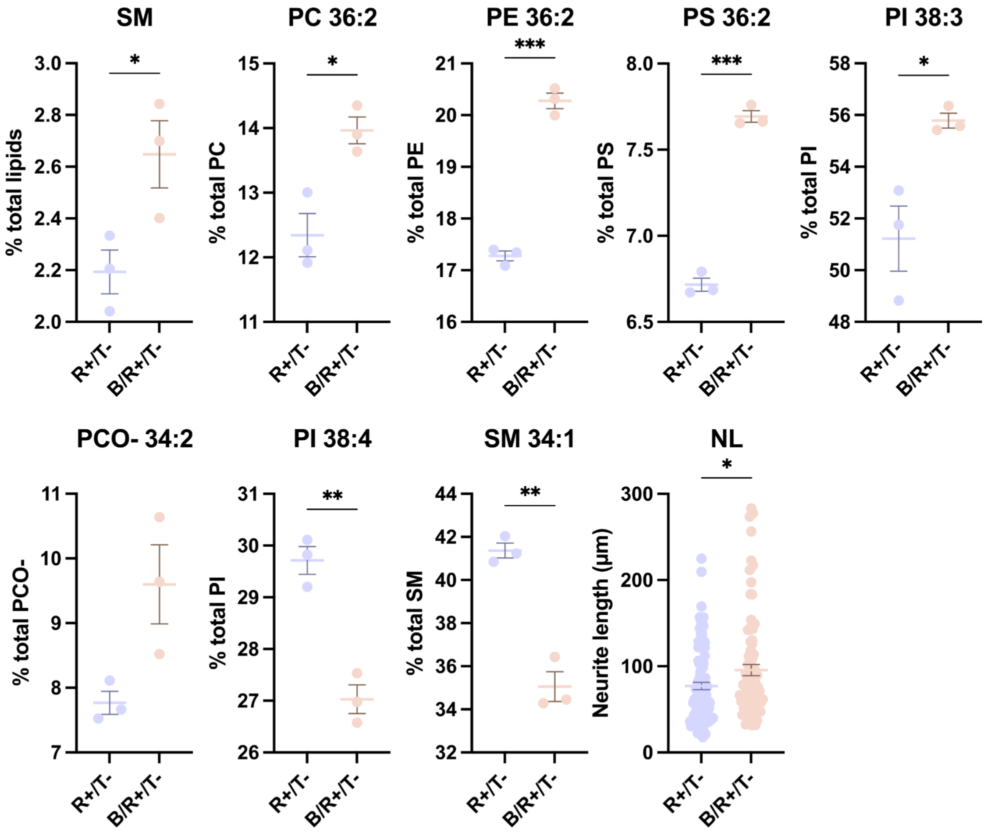


**Figure S3: Effect of BMP4 pre-treatment on the lipid levels and NL. t-test to compare means of R+/T- and B/R+/T-, *p<0.033, **p<0.002, ***p<0.001 (see Table S1 for p-values and statistical analysis details). Data shown as mean ± SEM and derive from 3 independent differentiations.**

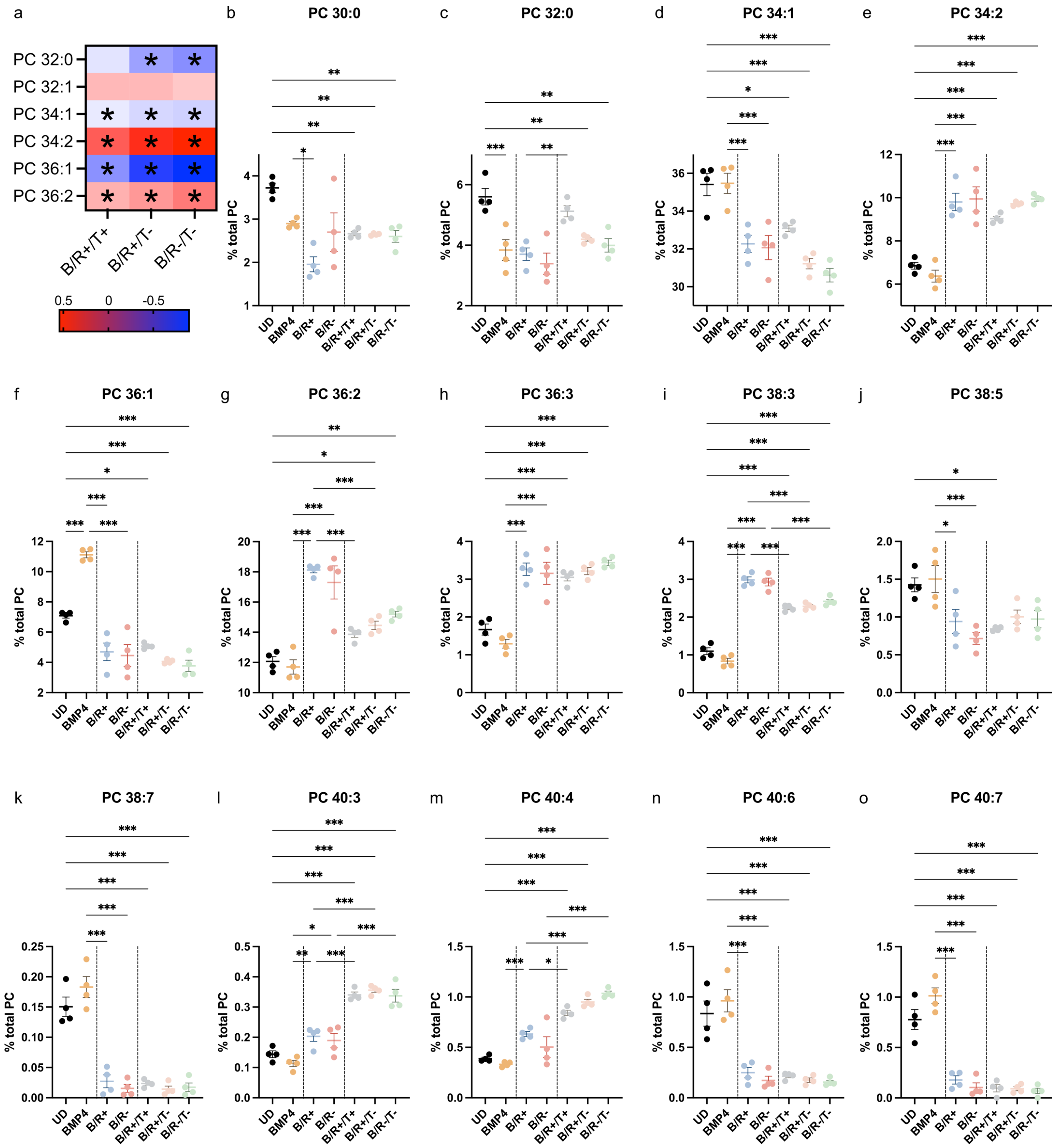


**Figure S4: The differentiation of SH-SY5Y cells leads to the significant changes in the levels of specific PC species. a. Heat map shows the FC (BRT/UD) in the levels of the most abundant species of PC (*for p<0.033, p<0.002, ***p<0.001, one-way ANOVA (UD, BMP4, B/R+, B/R-, B/R+/T+, B/R+/T-, B/R-/T-). b-o. Changes in the levels of PC species along the differentiation process. Only species whose levels are significantly affected by at least one BRT treatment and that have been detected in all UD and BRT replicates are shown. One-way ANOVA to compare means of UD, BMP4, B/R+, B/R-, B/R+/T+, B/R+/T- and B/R-/T-, *p<0.033, **p<0.002, ***p<0.001 (see Table S1 for p-values and statistical analysis details). Comparison shown: B/R+/T+, B/R+/T- and B/R-/T- vs UD (a-o), BMP4 vs UD (b-o), B/R+ and B/R- vs BMP4 (b-o), B/R+/T+, B/R+/T- and B/R-/T- vs their respective B/R(+/-) (b-o). Data shown as mean ± SEM and derive from 4 independent differentiations.**

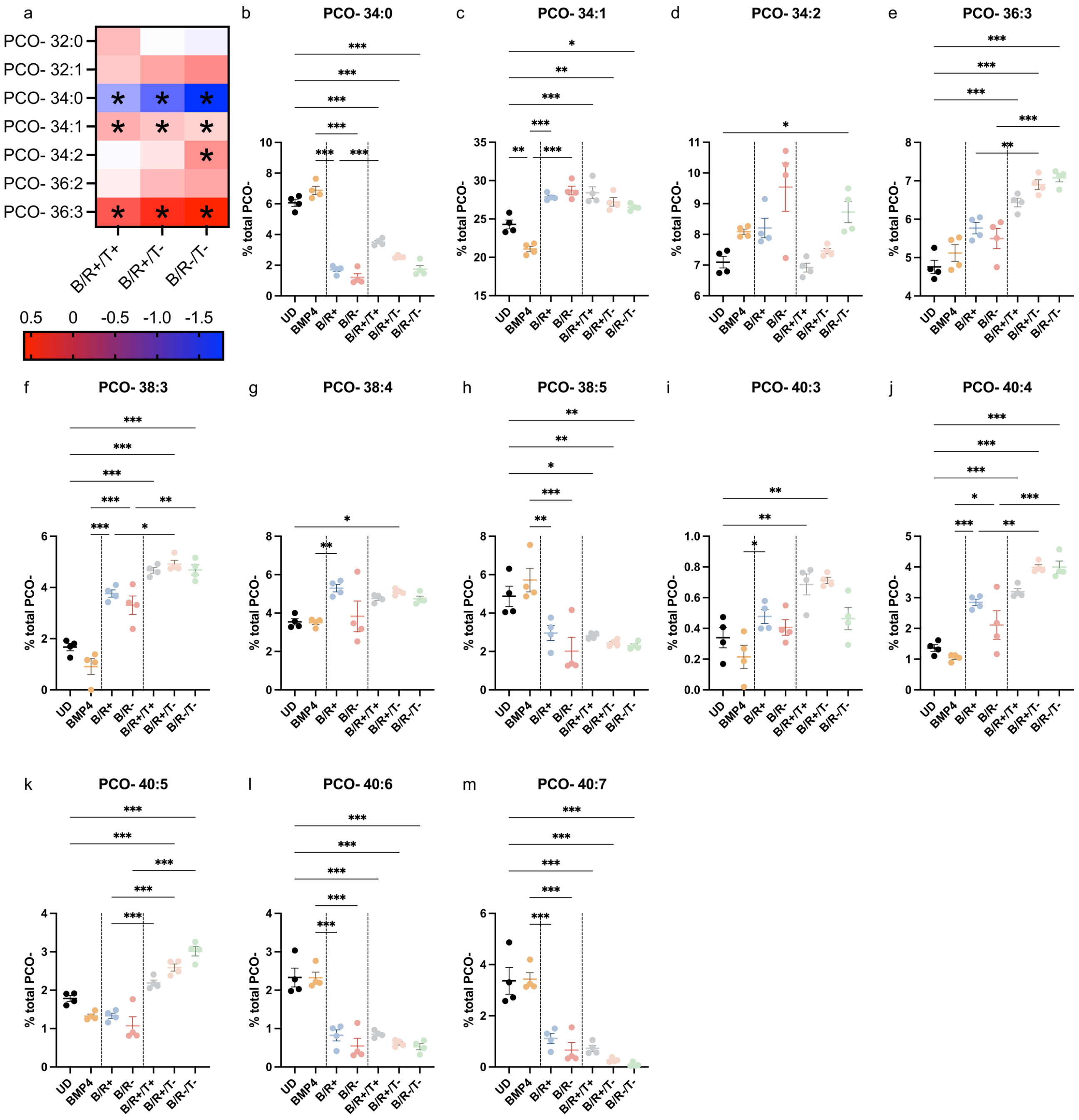


**Figure S5: The differentiation of SH-SY5Y cells leads to the significant changes in the levels of specific PCO- species. a. Heat map shows the FC (BRT/UD) in the levels of the most abundant species of PCO- (*for p<0.033, p<0.002, ***p<0.001, one-way ANOVA (UD, BMP4, B/R+, B/R-, B/R+/T+, B/R+/T-, B/R-/T-). b-m. Changes in the levels of PCO- species along the differentiation process. Only species whose levels are significantly affected by at least one BRT treatment and that have been detected in all UD and BRT replicates are shown. One-way ANOVA to compare means of UD, BMP4, B/R+, B/R-, B/R+/T+, B/R+/T- and B/R-/T-, *p<0.033, **p<0.002, ***p<0.001 (see Table S1 for p-values and statistical analysis details). Comparison shown: B/R+/T+, B/R+/T- and B/R-/T- vs UD (a-m), BMP4 vs UD (b-m), B/R+ and B/R- vs BMP4 (b-m), B/R+/T+, B/R+/T- and B/R-/T- vs their respective B/R(+/-) (b-m). Data shown as mean ± SEM and derive from 4 independent differentiations.**

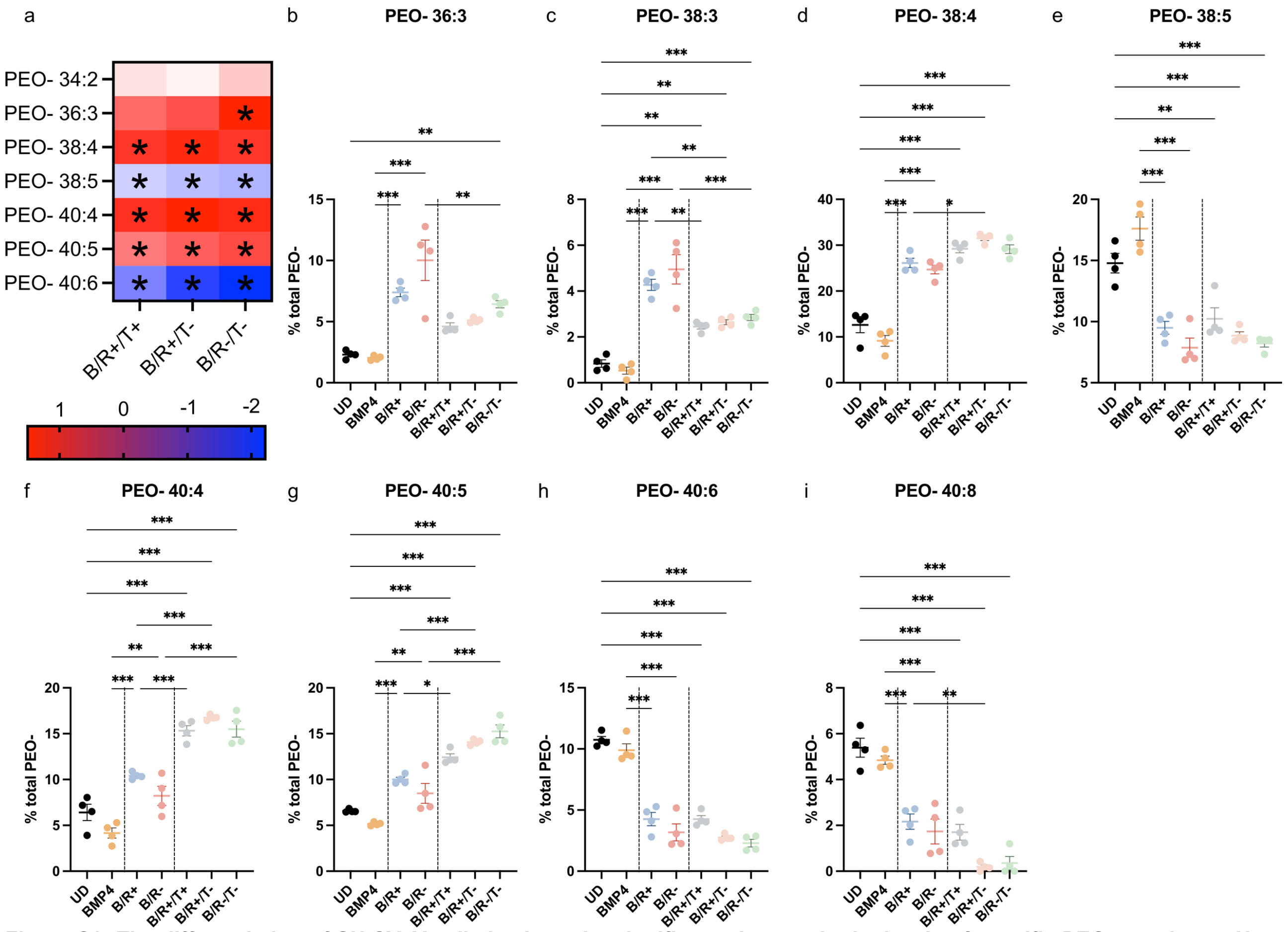


**Figure S6: The differentiation of SH-SY5Y cells leads to the significant changes in the levels of specific PEO- species. a. Heat map shows the FC (BRT/UD) in the levels of the most abundant species of PEO- (*for p<0.033, p<0.002, p<0.001, one-way ANOVA (UD, BMP4, B/R+, B/R-, B/R+/T+, B/R+/T-, B/R-/T-). b-i. Changes in the levels of PEO- species along the differentiation process. Only species whose levels are significantly affected by at least one BRT treatment and that have been detected in all UD and BRT replicates are shown. One-way ANOVA to compare means of UD, BMP4, B/R+, B/R-, B/R+/T+, B/R+/T- and B/R-/T-, *p<0.033, **p<0.002, ***p<0.001 (see Table S1 for p-values and statistical analysis details). Comparison shown: B/R+/T+, B/R+/T- and B/R-/T- vs UD (a-i), BMP4 vs UD (b-i), B/R+ and B/R- vs BMP4 (b-i), B/R+/T+, B/R+/T- and B/R-/T- vs their respective B/R(+/-) (b-i). Data shown as mean ± SEM and derive from 4 independent differentiations.**

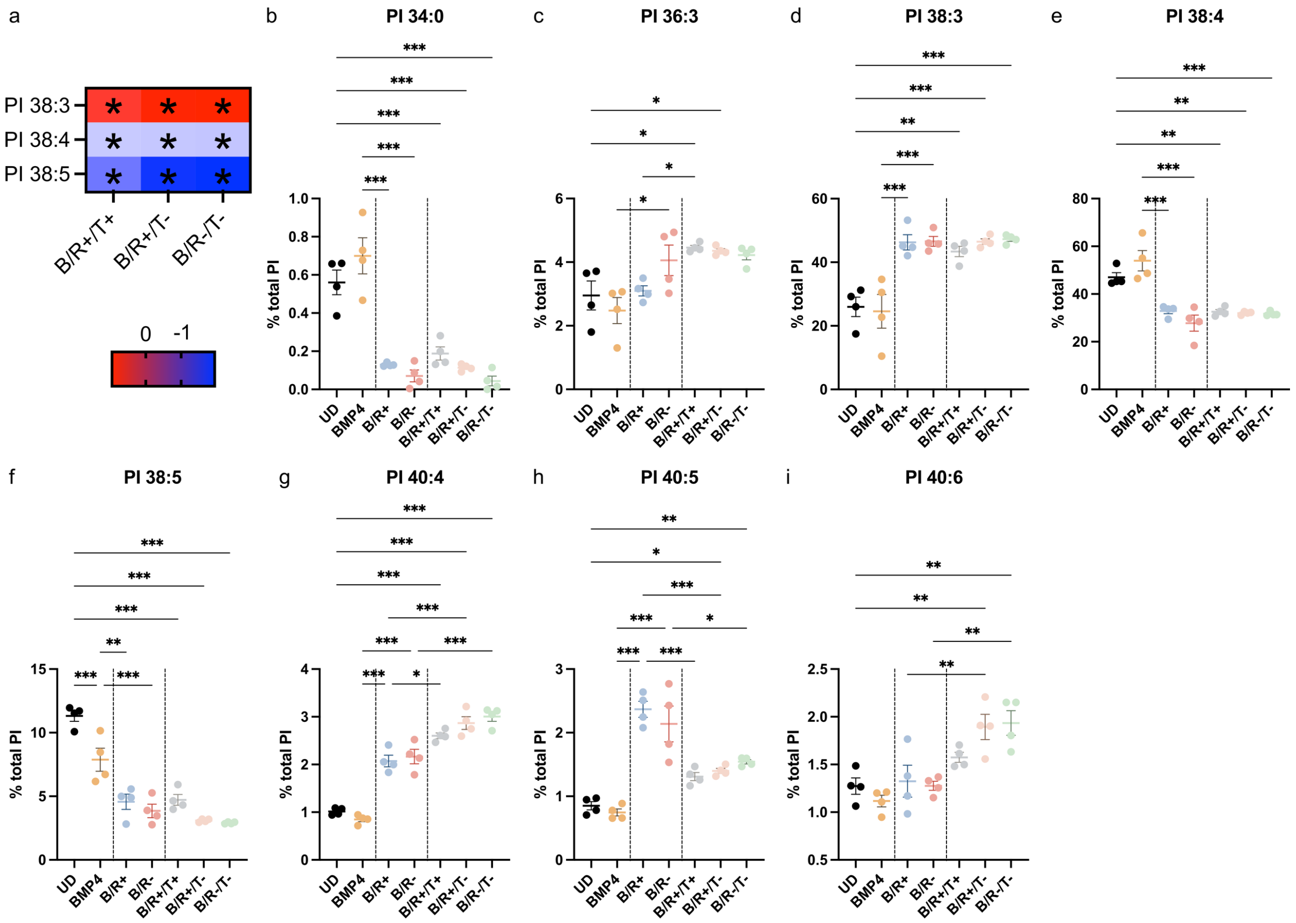


**Figure S7: The differentiation of SH-SY5Y cells leads to the significant changes in the levels of specific PI species. a. Heat map shows the FC (BRT/UD) in the levels of the most abundant species of PI (*for p<0.033, p<0.002, p<0.001, one-way ANOVA (UD, BMP4, B/R+, B/R-, B/R+/T+, B/R+/T-, B/R-/T-). b-i. Changes in the levels of PI species along the differentiation process. Only species whose levels are significantly affected by at least one BRT treatment and that have been detected in all UD and BRT replicates are shown. One-way ANOVA to compare means of UD, BMP4, B/R+, B/R-, B/R+/T+, B/R+/T- and B/R-/T-, *p<0.033, **p<0.002, ***p<0.001 (see Table S1 for p-values and statistical analysis details). Comparison shown: B/R+/T+, B/R+/T- and B/R-/T- vs UD (a-i), BMP4 vs UD (b-i), B/R+ and B/R- vs BMP4 (b-i), B/R+/T+, B/R+/T- and B/R-/T- vs their respective B/R(+/-) (b-i). Data shown as mean ± SEM and derive from 4 independent differentiations.**

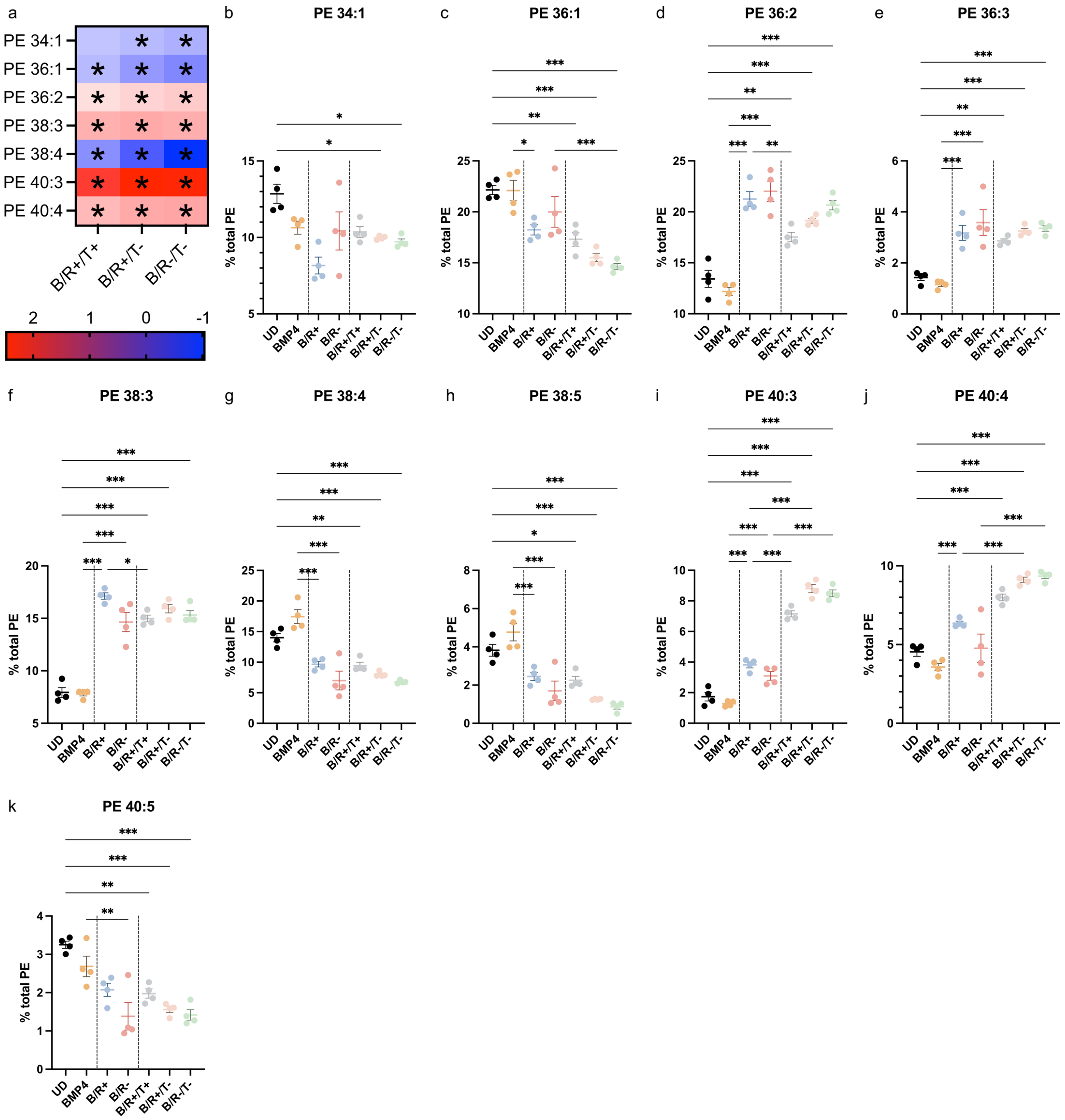

**Figure S8: The differentiation of SH-SY5Y cells leads to the significant changes in the levels of specific PE species. a. Heat map shows the FC (BRT/UD) in the levels of the most abundant species of PE (*for p<0.033, p<0.002, p<0.001, one-way ANOVA (UD, BMP4, B/R+, B/R-, B/R+/T+, B/R+/T-, B/R-/T-). b-k. Changes in the levels of PE species along the differentiation process. Only species whose levels are significantly affected by at least one BRT treatment and that have been detected in all UD and BRT replicates are shown. One-way ANOVA to compare means of UD, BMP4, B/R+, B/R-, B/R+/T+, B/R+/T- and B/R-/T-, *p<0.033, **p<0.002, ***p<0.001 (see Table S1 for p-values and statistical analysis details). Comparison shown: B/R+/T+, B/R+/T- and B/R-/T- vs UD (a-k), BMP4 vs UD (b-k), B/R+ and B/R- vs BMP4 (b-k), B/R+/T+, B/R+/T- and B/R-/T- vs their respective B/R(+/-) (b-k). Data shown as mean ± SEM and derive from 4 independent differentiations.**

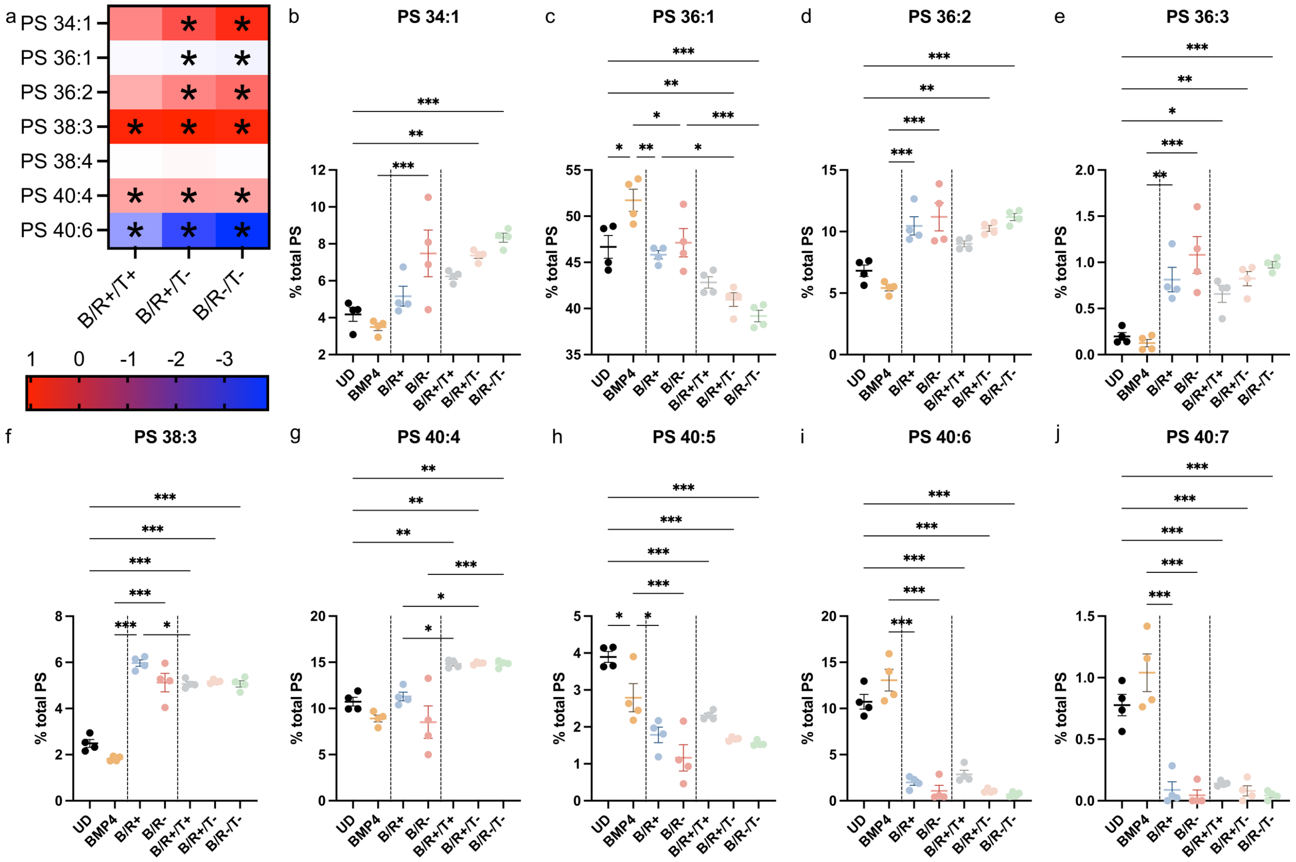


**Figure S9: The differentiation of SH-SY5Y cells leads to the significant changes in the levels of specific PS species. a. Heat map shows the FC (BRT/UD) in the levels of the most abundant species of PS (*for p<0.033, p<0.002, p<0.001, one-way ANOVA (UD, BMP4, B/R+, B/R-, B/R+/T+, B/R+/T-, B/R-/T-). b-j. Changes in the levels of PS species along the differentiation process. Only species whose levels are significantly affected by at least one BRT treatment and that have been detected in all UD and BRT replicates are shown. One-way ANOVA to compare means of UD, BMP4, B/R+, B/R-, B/R+/T+, B/R+/T- and B/R-/T-, *p<0.033, **p<0.002, ***p<0.001 (see Table S1 for p-values and statistical analysis details). Comparison shown: B/R+/T+, B/R+/T- and B/R-/T- vs UD (a-j), BMP4 vs UD (b-j), B/R+ and B/R- vs BMP4 (b-j), B/R+/T+, B/R+/T- and B/R-/T- vs their respective B/R(+/-) (b-j). Data shown as mean ± SEM and derive from 4 independent differentiations.**

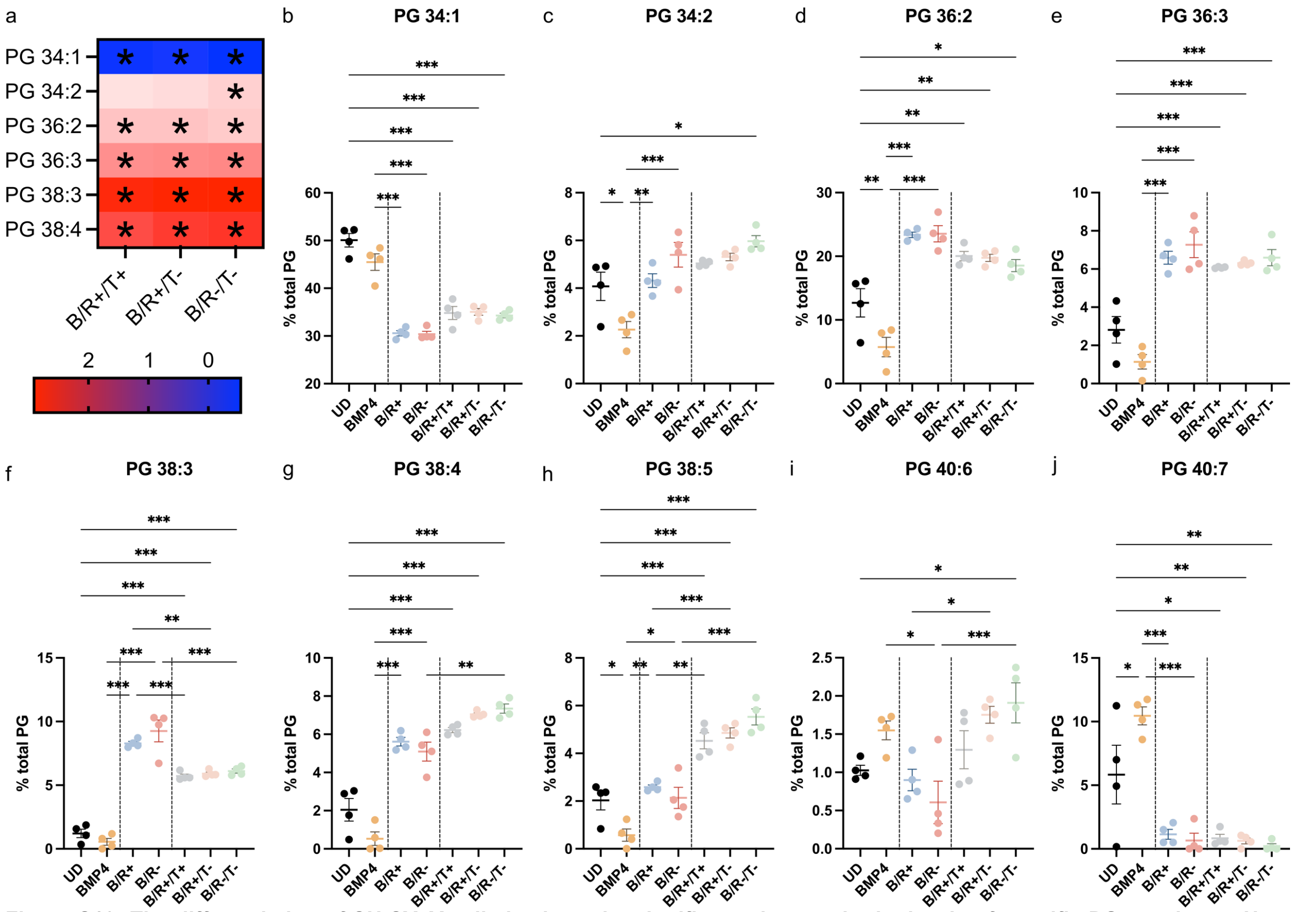

**Figure S10: The differentiation of SH-SY5Y cells leads to the significant changes in the levels of specific PG species. a. Heat map shows the FC (BRT/UD) in the levels of the most abundant species of PG (*p<0.033, p<0.002, p<0.001, one-way ANOVA (UD, BMP4, B/R+, B/R-, B/R+/T+, B/R+/T-, B/R-/T-). b-j. Changes in the levels of PG species along the differentiation process. Only species whose levels are significantly affected by at least one BRT treatment and that have been detected in all UD and BRT replicates are shown. One-way ANOVA to compare means of UD, BMP4, B/R+, B/R-, B/R+/T+, B/R+/T- and B/R-/T-, *p<0.033, **p<0.002, ***p<0.001 (see Table S1 for p-values and statistical analysis details). Comparison shown: B/R+/T+, B/R+/T- and B/R-/T- vs UD (a-j), BMP4 vs UD (b-j), B/R+ and B/R- vs BMP4 (b-j), B/R+/T+, B/R+/T- and B/R-/T- vs their respective B/R(+/-) (b-j). Data shown as mean ± SEM and derive from 4 independent differentiations.**

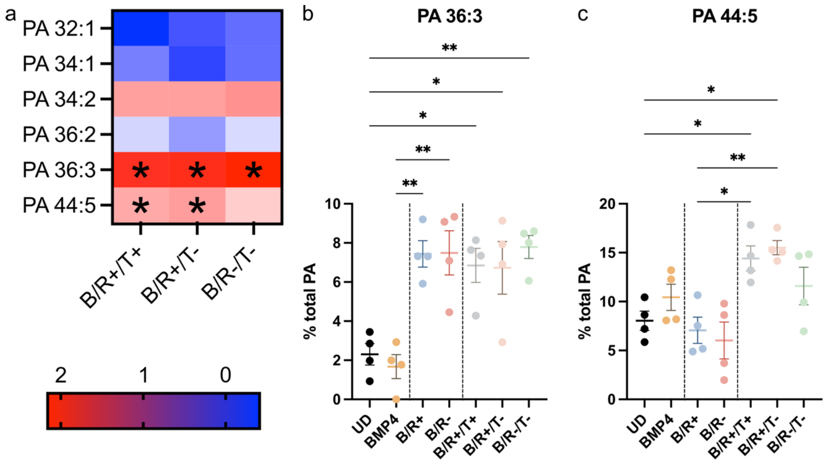


**Figure S11: The differentiation of SH-SY5Y cells leads to the significant changes in the levels of specific PA species. a. Heat map shows the FC (BRT/UD) in the levels of the most abundant species of PA (*p<0.033, p<0.002, p<0.001, one-way ANOVA (UD, BMP4, B/R+, B/R-, B/R+/T+, B/R+/T-, B/R-/T-). b-c. Changes in the levels of PA species along the differentiation process. Only species whose levels are significantly affected by at least one BRT treatment and that have been detected in all UD and BRT replicates are shown. One-way ANOVA to compare means of UD, BMP4, B/R+, B/R-, B/R+/T+, B/R+/T- and B/R-/T-, *p<0.033, **p<0.002, ***p<0.001 (see Table S1 for p-values and statistical analysis details). Comparison shown: B/R+/T+, B/R+/T- and B/R-/T- vs UD (a-c), BMP4 vs UD (b-c), B/R+ and B/R- vs BMP4 (b-c), B/R+/T+, B/R+/T- and B/R-/T- vs their respective B/R(+/-) (b-c). Data shown as mean ± SEM and derive from 4 independent differentiations.**

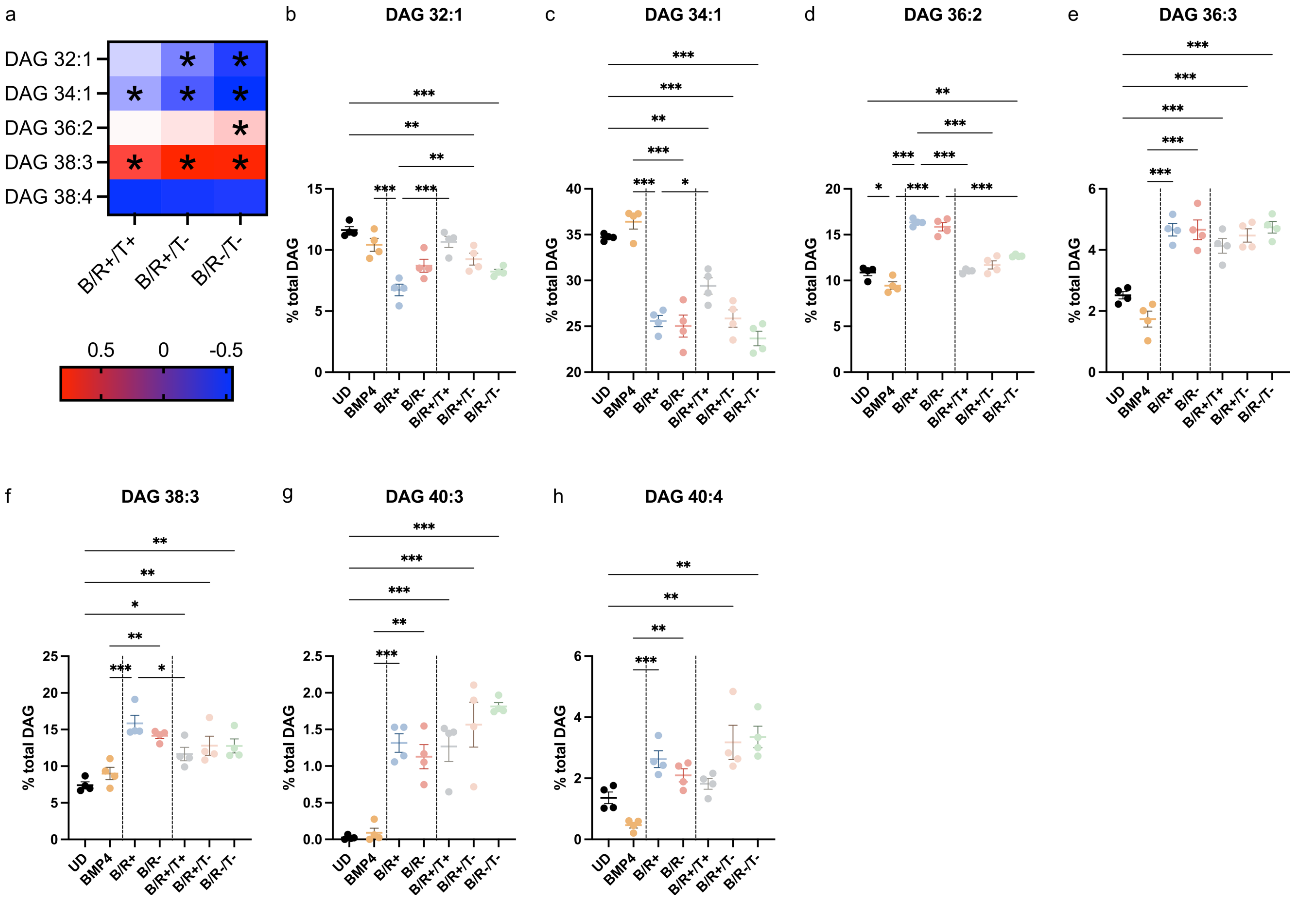


**Figure S12: The differentiation of SH-SY5Y cells leads to the significant changes in the levels of specific DAG species. a. Heat map shows the FC (BRT/UD) in the levels of the most abundant species of DAG (*p<0.033, p<0.002, p<0.001, one-way ANOVA (UD, BMP4, B/R+, B/R-, B/R+/T+, B/R+/T-, B/R-/T-). b-h. Changes in the levels of DAG species along the differentiation process. Only species whose levels are significantly affected by at least one BRT treatment and that have been detected in all UD and BRT replicates are shown. One-way ANOVA to compare means of UD, BMP4, B/R+, B/R-, B/R+/T+, B/R+/T- and B/R-/T-, *p<0.033, **p<0.002, ***p<0.001 (see Table S1 for p-values and statistical analysis details). Comparison shown: B/R+/T+, B/R+/T- and B/R-/T- vs UD (a-h), BMP4 vs UD (b-h), B/R+ and B/R- vs BMP4 (b-h), B/R+/T+, B/R+/T- and B/R-/T- vs their respective B/R(+/-) (b-h). Data shown as mean ± SEM and derive from 4 independent differentiations.**

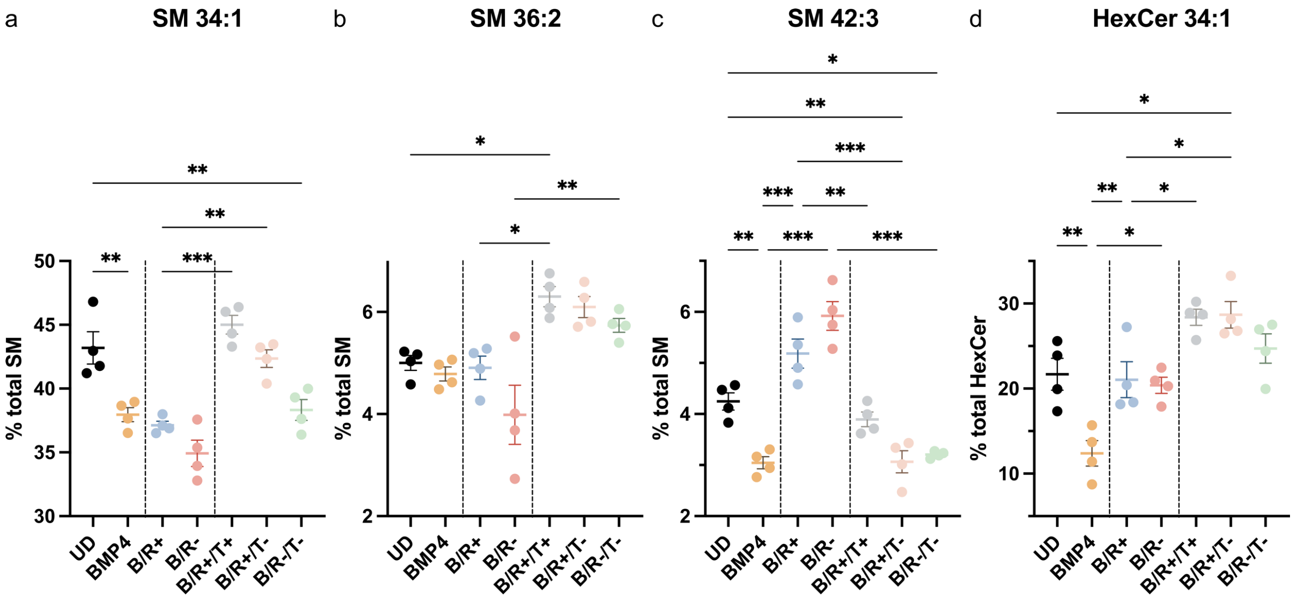


**Figure S13: The differentiation of SH-SY5Y cells leads to the significant changes in the levels of SL species. a-d. Changes in the levels of SL species along the differentiation process. Only species whose levels are significantly affected by at least one BRT treatment and that have been detected in all UD and BRT replicates are shown. One-way ANOVA to compare means of UD, BMP4, B/R+, B/R-, B/R+/T+, B/R+/T- and B/R-/T-, *p<0.033, **p<0.002, ***p<0.001 (see Table S1 for p-values and statistical analysis details). Comparison shown: B/R+/T+, B/R+/T- and B/R-/T- vs UD, BMP4 vs UD, B/R+ and B/R- vs BMP4, B/R+/T+, B/R+/T- and B/R-/T- vs their respective B/R(+/-). Data shown as mean ± SEM and derive from 4 independent differentiations.**

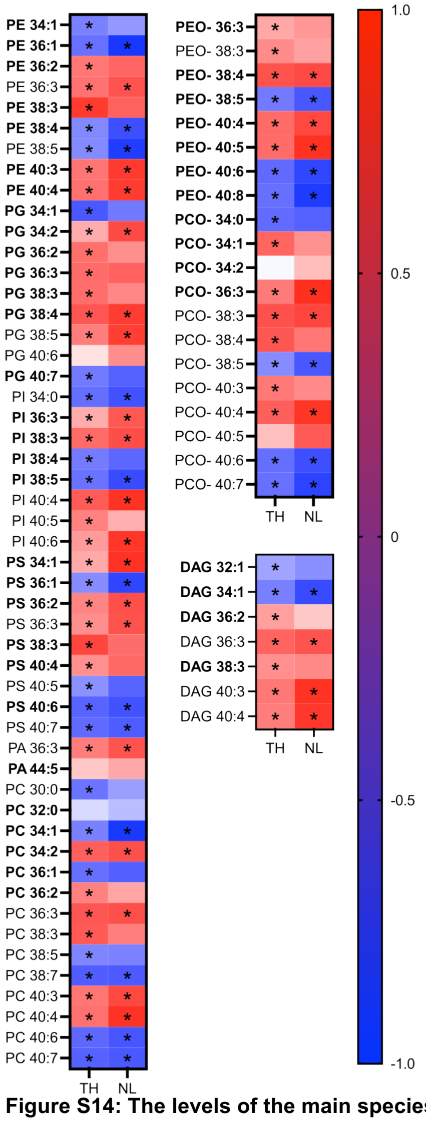


**Figure S14: The levels of the main species of phospholipids, ether phospholipids and diacylglycerides correlate with TH levels and/or neurite length. Heat maps representing Pearson r for the correlation between the percentage of phospholipid species vs OD (TH/β-actin) (TH) or neurite length for the data sets UD, BMP4, B/R+, B/R-, B/R+/T+, B/R+/T-, B/R-/T- (*p<0.033) (see Table S1 for p-values and statistical analysis details). The main species for each lipid class are shown in bold.**

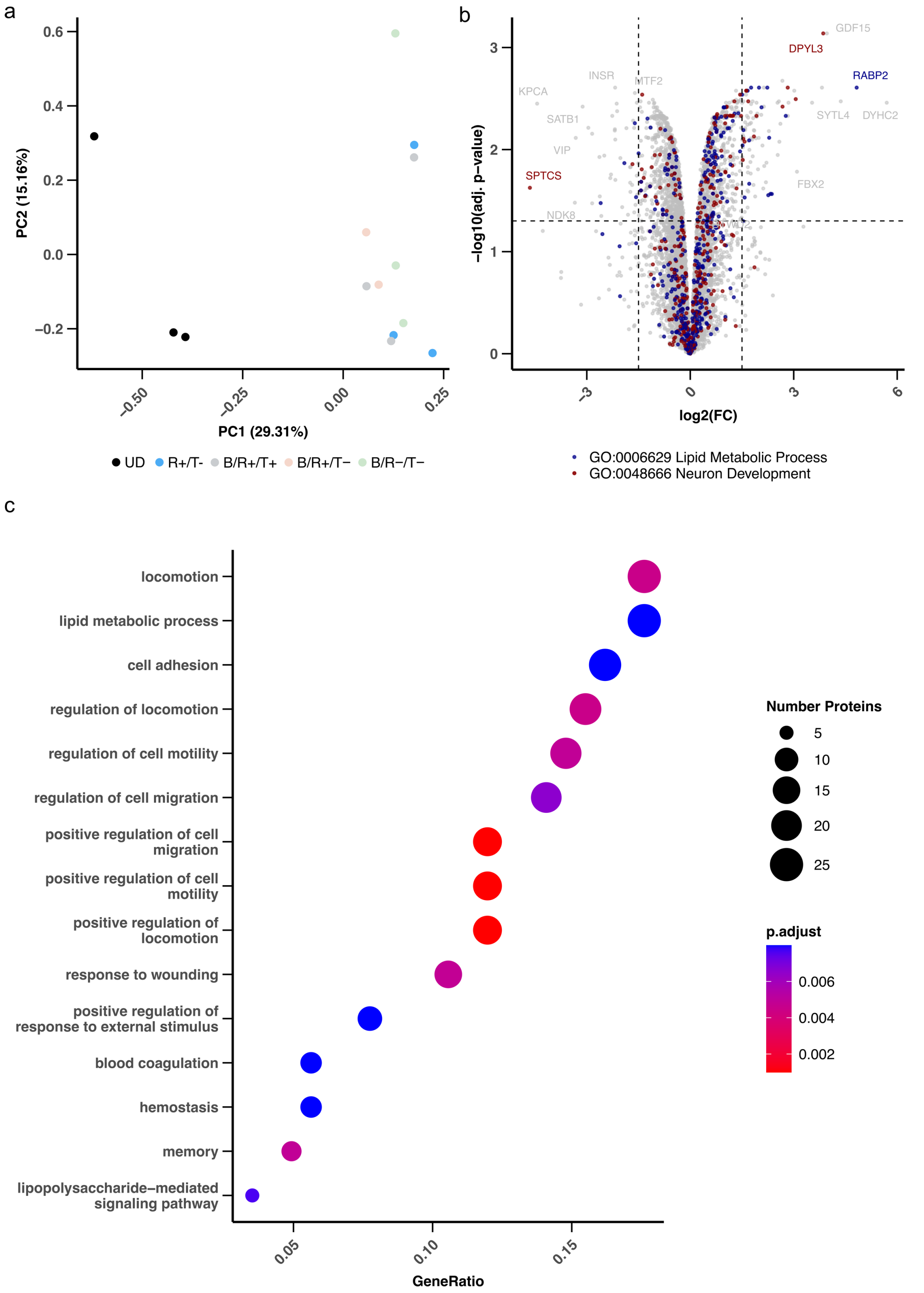


**Figure S15: BRT-induced differentiation of SH-SY5Y cells leads to changes in the levels of proteins involved in lipid metabolism and neuron maturation. (a) Principal component analysis of protein expression of 6071 quantified proteins in all replicates. No data imputation. (b) Differentially expressed proteins between undifferentiated and B/R+/T- treated SH-SY5Y cells. (c) GO over-representation analysis of the differentially expressed proteins between undifferentiated and B/R+/T- treated SH-SY5Y cells. The 15 most over-representation biological function GO terms are shown with the number of proteins related to them, the adjusted p-value and the gene ratio.**

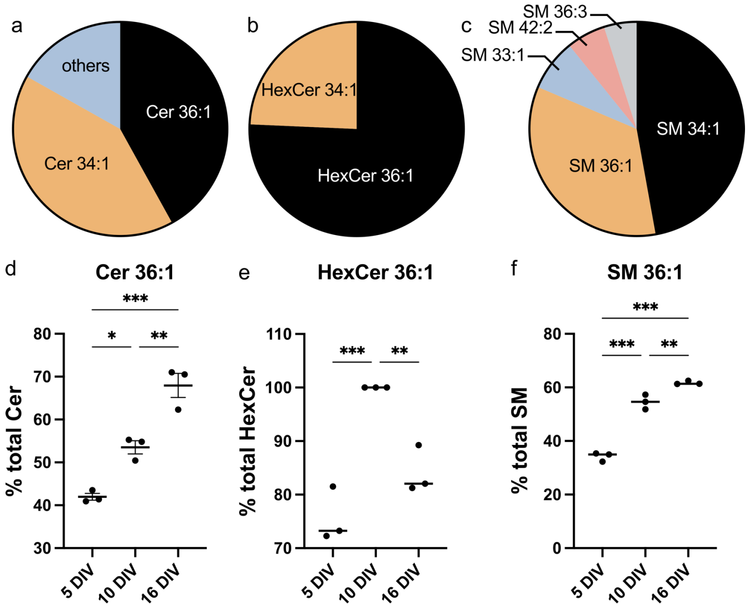


**Figure S16: The maturation of primary neurons leads to an increase in the 36:1 species of Cer, HexCer and SM. a-c. Distribution of the species of Cer, HexCer and SM in primary neurons (data from Fitzner et al). d-f. Levels of Cer 36:1 (d), HexCer 36:1 (e) and SM 36:1 (f) in primary neurons after 5, 10 and 16 days in vitro (DIV) (data from Fitzner et al). One-way ANOVA to compare means of 5 DIV, 10 DIV, 16 DIV, *p<0.033, **p<0.002, ***p<0.001 (see Table S1 for p-values and statistical analysis details). Data shown as mean ± SEM.**
